# Viral entry is a weak barrier to zoonosis

**DOI:** 10.1101/2024.01.22.576693

**Authors:** Jérémy Dufloo, Iván Andreu-Moreno, Ana Valero-Rello, Rafael Sanjuán

## Abstract

Recent advances in viral metagenomics have led to the discovery of many mammalian viruses, but experimental tests to determine whether they pose a threat to humans are largely lacking. A first step for a virus to cross the species barrier is to penetrate host cells. Here, we use gene synthesis and viral pseudotyping to experimentally test the ability of viral receptor-binding proteins (RBPs) from >100 enveloped RNA viruses to mediate entry into human cells. Analysis of thousands of RBP-cell pairs demonstrated such ability for most viruses, with significant variation among the 14 viral families studied. Comparison of RBP-mediated infectivity with cellular gene expression data showed that viral entry is often not limited by the presence of a receptor and revealed the contribution of additional host factors. Our results prove the weakness of interspecies barriers at the early stages of infection and identify molecular interactions that enable viral zoonosis.

## Introduction

The cross-species transmission of animal viruses to humans (zoonosis) poses a tremendous threat to human health, as exemplified by pandemics such as COVID-19, AIDS, and the Spanish flu. A substantial fraction of the estimated 40,000 viruses infecting wildlife or domestic mammals may spill over into humans^1^. However, predicting which viruses are more likely to emerge is extremely challenging, since this is a largely random process influenced by many ecological, social, genetic, and virological factors. Previous studies have identified ecological risk factors including biodiversity loss^2^, global warming^3^, and other disturbances^4,5^. Certain viral traits have also been associated with zoonotic risk. In particular, enveloped RNA viruses are of greatest concern as they exhibit increased cross-species transmissibility and account for >70% of all zoonotic viruses^6–9^.

Characterizing the animal virome is the first step towards the identification of zoonotic threats, and recent large-scale viral metagenomics initiatives have led to major advances in this area^10^. In contrast, experimental virology studies providing functional information on these new viruses are scarce^11^. Whether wildlife viruses can infect human cells is usually unknown because viral culturing is technically challenging and raises biosafety issues, hindering its large-scale applicability. One way to circumvent these limitations is to use surrogate systems that recapitulate specific steps of the infection cycle, such as viral pseudotypes, in which the receptor-binding protein (RBP) of an enveloped virus of interest is incorporated into a viral vector. Pseudotyping has been applied to most families of enveloped RNA viruses and can faithfully reproduce key processes such as receptor usage, cellular tropism, viral entry routes, and antibody-mediated neutralization^12^.

The cross-species transmission of viruses initially depends on the compatibility between the viral RBP and the cellular entry factors of different host species. Interspecies variability in these factors is often considered a major barrier to zoonosis, based on observations made with a few well-studied viruses, such as avian and human influenza strains^13^ and coronaviruses^14^ among others. Moreover, an evolutionary arms race between RBPs and specific virus receptors has been demonstrated in several cases including some rodent arenaviruses^15^, bat ebolaviruses^16^, and hominid lentiviruses^17^. However, viral entry is a complex process that typically involves multiple steps including initial attachment^18^, receptor binding, endocytosis pathways, and antiviral proteins acting at the entry stage of infection^19^. How these factors determine the infection of human cells remains unknown for most viruses.

Since many RBP sequences from uncultured viruses infecting wild and domestic mammals are now available, the combination of gene synthesis and pseudotyping allows us to systematically study the human transmissibility of uncharacterized viruses at the entry stage of infection, and to investigate the molecular determinants of this process. To achieve this goal, we engineered pseudotypes carrying the RBPs from over a hundred viral species belonging to different families of enveloped RNA viruses and tested them in dozens of well-characterized human cell lines. We detected viral entry for most RBP-cell combinations. However, infectivity varied strongly across viral families, with coronaviruses, flaviviruses, and matonaviruses showing the strongest interspecies barrier at the entry stage. We also found that specific RBP-receptor interactions are not necessarily a limiting step for viral infection, and we identified host factors with broad-range effects on viral entry and cellular tropism.

## Results

### Viral entry is a weak barrier to zoonosis

We used vesicular stomatitis virus (VSV) as a vector for the production of viral pseudotypes carrying heterologous RBPs (**Figure S1A**). For this, we first built RBP phylogenetic trees for 14 different families of enveloped RNA viruses (**Figures S2-S15)**, from which we selected 126 representative RBPs and obtained their sequences through gene synthesis. We successfully constructed VSV pseudotypes for 102 of these RBPs, as shown by infectivity tests or detection of RBP incorporation into viral particles by Western blot (**Figures S2-S15**). Of these, 79 corresponded to viruses not reported to infect humans to date, including 56 viruses described in mammals such as bats, rodents, or artiodactyls, among others, 5 to viruses described in other vertebrates, and 18 to viruses discovered in arthropods but belonging to families that typically infect mammals (**Table S1**). Of the 56 mammalian viruses, 41 have only been found in wildlife and 15 infect domestic mammals such as cattle, camels, or cats. As controls, we also included 20 known zoonotic viruses as well as 3 human-exclusive viruses.

To preliminarily test whether the 102 RBPs could mediate viral entry into human cells, we inoculated the widely used HEK293T cell line with each pseudotype. Infection was observed in 74 cases (**Figure S16**), revealing a weak interspecies barrier at the level of viral entry. To examine this more thoroughly, we set out to extend our experiments to 48 human cell lines from the NCI-60 reference panel (**Table S2**). We verified that the overall expression of cell surface genes in these cell lines correlated with the values reported for normal tissues in the Human Protein Atlas (Pearson r = 0.840, P < 0.001; **Figure S17A**), and that 48 cell lines was a sufficient sample size to detect human-tropic RBPs (**Figure S17B**). Out of the 102 × 48 = 4896 total RBP-cell pairs tested, 2675 (54.6%) reproducibly showed infection (**Figure 1; Table S3**), and 82/102 (80.4%) pseudotypes infected at least one of the 48 cell lines. Thus, the availability of suitable entry factors does not constitute a major obstacle to viral zoonosis.

**Figure 1.**
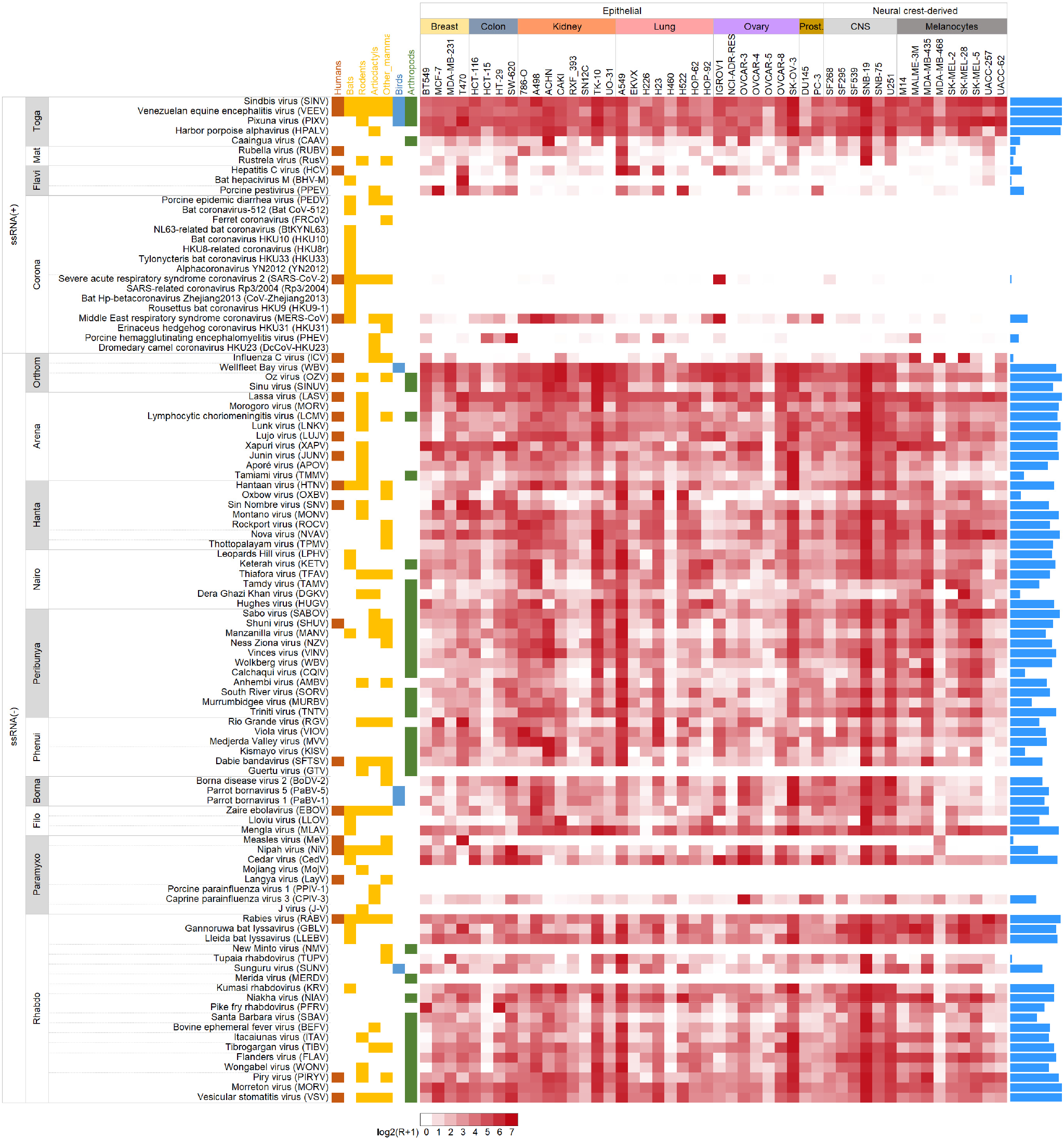
Human cell tropism for the RBPs of 102 enveloped RNA viruses. Viruses are sorted by family and organized by genera within families, as indicated by the horizontal lines. Yellow, orange and green boxes show whether each virus has been reported to infect non-human mammals, humans, and arthropods, respectively. Names of the 48 cell lines are indicated at the top, and cells are organized by tissue of origin. The heat map shows relative pseudotype infectivity, calculated as log_2_(R+1), where R is the multiplicity of infection (MOI) scaled as a percentage of the maximum MOI across the 48 cell lines for each pseudotype. Blue bars on the right indicate the number of cell lines in which infection was detected. Data are provided in **Table S2**.

### RBP human cell tropism varies according to viral and host traits

We found large variations in human cell entry among RBPs belonging to different viral families (logistic regression, P < 0.001). Coronavirus, matonavirus, and flavivirus pseudotypes exhibited the narrowest tropism, infecting only 3.3%, 8.3%, and 17.4% of the 48 cell lines, respectively (**Figure 2A**). Indeed, coronaviruses comprised 13 of the 20 pseudotypes showing a complete lack of infectivity. Moreover, the three coronavirus pseudotypes for which we detected infection entered only 1, 8, and 16 of the 48 cell lines. For instance, the SARS-CoV-2 spike mediated entry into IGROV-1 cells, as shown previously^20^, but none of the other 47 cell lines gave any infection signal above the background level. In contrast, pseudotypes from the other 11 viral families infected most of the cell lines, with the highest percentages corresponding to arenaviruses (84.50 ± 7.6%), togaviruses (83.3 ± 16.2%), and hantaviruses (75.9 ± 9.6%).

**Figure 2.**
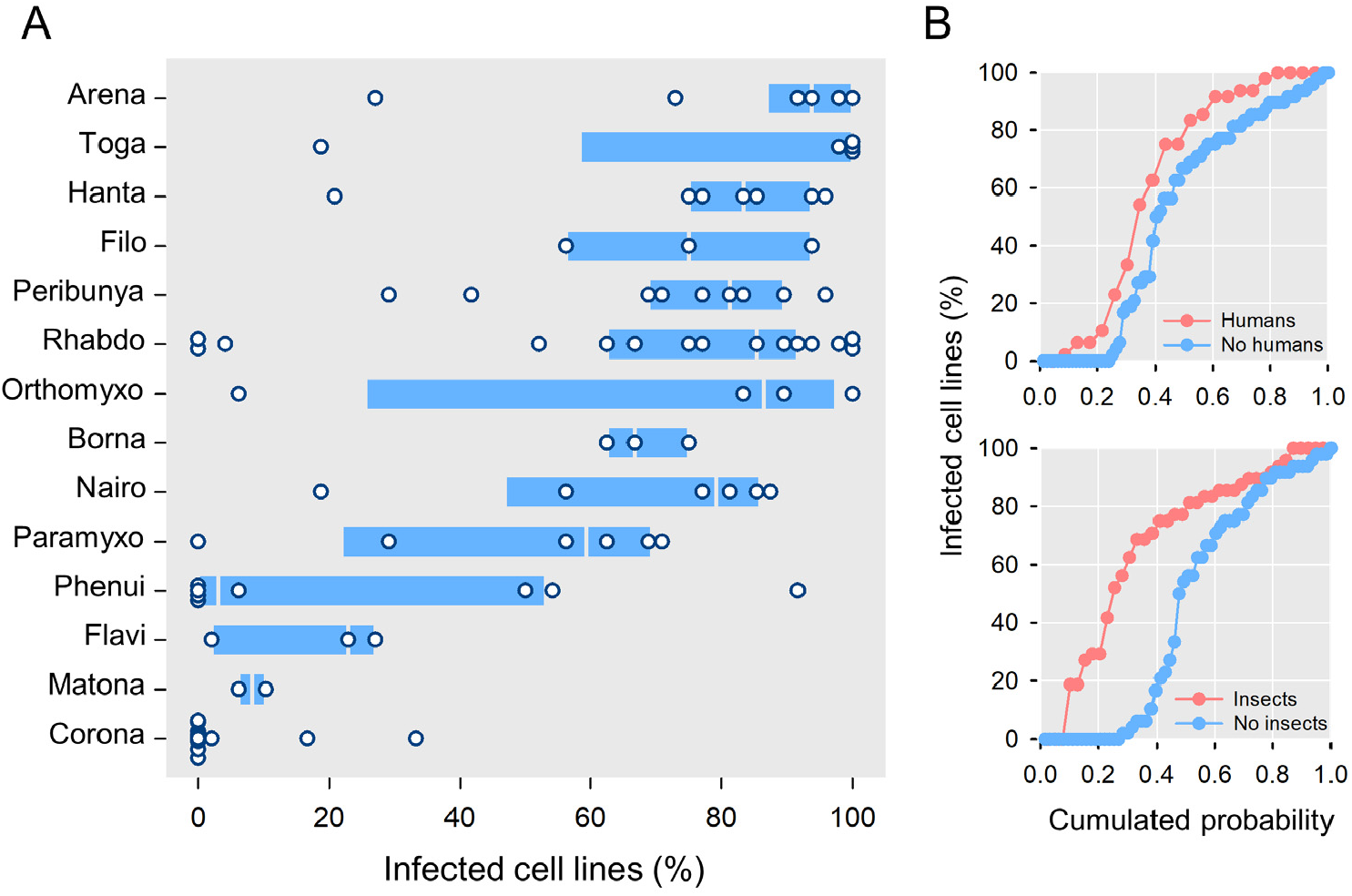
Association of viral and host traits with RBP cell tropism. **A**. Variation in the proportion of infected human cell lines according to viral family. The percentage of 48 lines for which infection was detected is shown. Boxes show the median (white line), 25^th^ and 75^th^ percentiles, and dots show data points for individual viruses. **B**. Cumulative probability distribution of the fraction of cell lines infected by the 102 pseudotypes as a function of whether viruses are known to infect humans (top) and whether they have been found in insects (bottom).

As expected, RBPs from viruses known to infect humans showed evidence of entry in a higher percentage of RBP-cell pairs tested than those from viruses not reported in humans. However, this difference was small (64.6 ± 8.0% versus 51.7 ± 4.2%; **Figure 2B**) and significant only for coronaviruses and rhabdoviruses (Wilcoxon rank-sum tests: P < 0.01). The lack of a detectable interspecies barrier at the entry stage was evident for the RBPs of most viruses described in rodents, bats, and other mammals. One interesting example was Rustrela virus, a relative of Rubella virus discovered in 2020 as the causative agent of brain infection in wild yellow-necked field mice and zoo animals^21^, for which we found RBP-mediated entry into human astrocyte- and lung-derived cells. A broad human cell tropism was observed even for the RBP of the Harbor porpoise alphavirus isolated in 2021 in Alaska from cetaceans^22^. Many pseudotypes from viruses discovered in non-mammalian hosts also infected most cell lines. Examples include Sinu virus, an orthomyxovirus isolated in 2017 from mosquitoes in Colombia^23^, Wellfleet Bay virus, another orthomyxovirus identified in 2014 as the causative agent of avian mass mortality in United States^24^, and Niakha virus, a rhabdovirus isolated in 2013 from phlebotomine sandflies in Senegal^25^. Indeed, pseudotypes of viruses infecting arthropods were particularly likely to infect human cells (Wilcoxon rank-sum test: P = 0.002; **Figure 2B**), demonstrating the extremely wide host range of these RBPs.

RBPs from known human-infective viruses also showed ample variation in their ability to enter different cell lines. Viruses with a broader cell tropism should be more likely to cause systemic acute infections and be highly pathogenic, as shown for well-studied viruses such as influenza^26^. One of the most infectious pseudotypes carried the hemagglutinin protein of Oz virus, a novel tick-borne zoonotic orthomyxovirus that caused the first human fatality in Japan in June 2023^27^. Infection rates >50% were also observed for the pseudotypes of Shuni orthobunyavirus, a suspected cause of human neurological disease in Africa^28^, Manzanilla orthobunyavirus, which is maintained in nature through a pig-mosquito-bird cycle in Africa and South East Asia^29^, Dabie bandavirus, a highly-pathogenic emerging bunyavirus responsible for the severe fever with thrombocytopenia syndrome in East Asia^30^, as well as several arenaviruses and rhabdoviruses.

We carried out a hierarchical clustering analysis to identify groups of RBPs showing similar cell tropisms. This yielded 20 clusters of 2-7 RBPs formed exclusively by members of the same family (**Figure 3**; **Figure S18**). Phylogenetically consistent clusters were found among orthobornaviruses, vesiculoviruses, quaranjaviruses, henipaviruses, bandaviruses, matonaviruses, and most peribunyaviruses, suggesting that viruses within these groups use similar entry pathways. However, in most cases, clusters did not include all members of a given taxonomical group. Besides viral phylogeny, we identified preference towards cells derived from the neuroectoderm as another factor driving that RBP clustering. This tropism was particularly marked in a well-delimited cluster formed by 8 peribunyavirus, 7 rhabodvirus, and 4 hantavirus RBPs (**Figure S19;** Wilcoxon rank-sum test: P < 0.001), including those from viruses known to cause brain pathology, such as Gannoruwa bat lyssavirus and rabies virus. In contrast, coronavirus and flavivirus pseudotypes rarely entered neuroectoderm-derived cells (0.4% and 0.0% cell lines, respectively), whereas epithelial cells were more frequently infected (4.5% and 25.32%, respectively; Fisher exact tests: P < 0.001).

**Figure 3.**
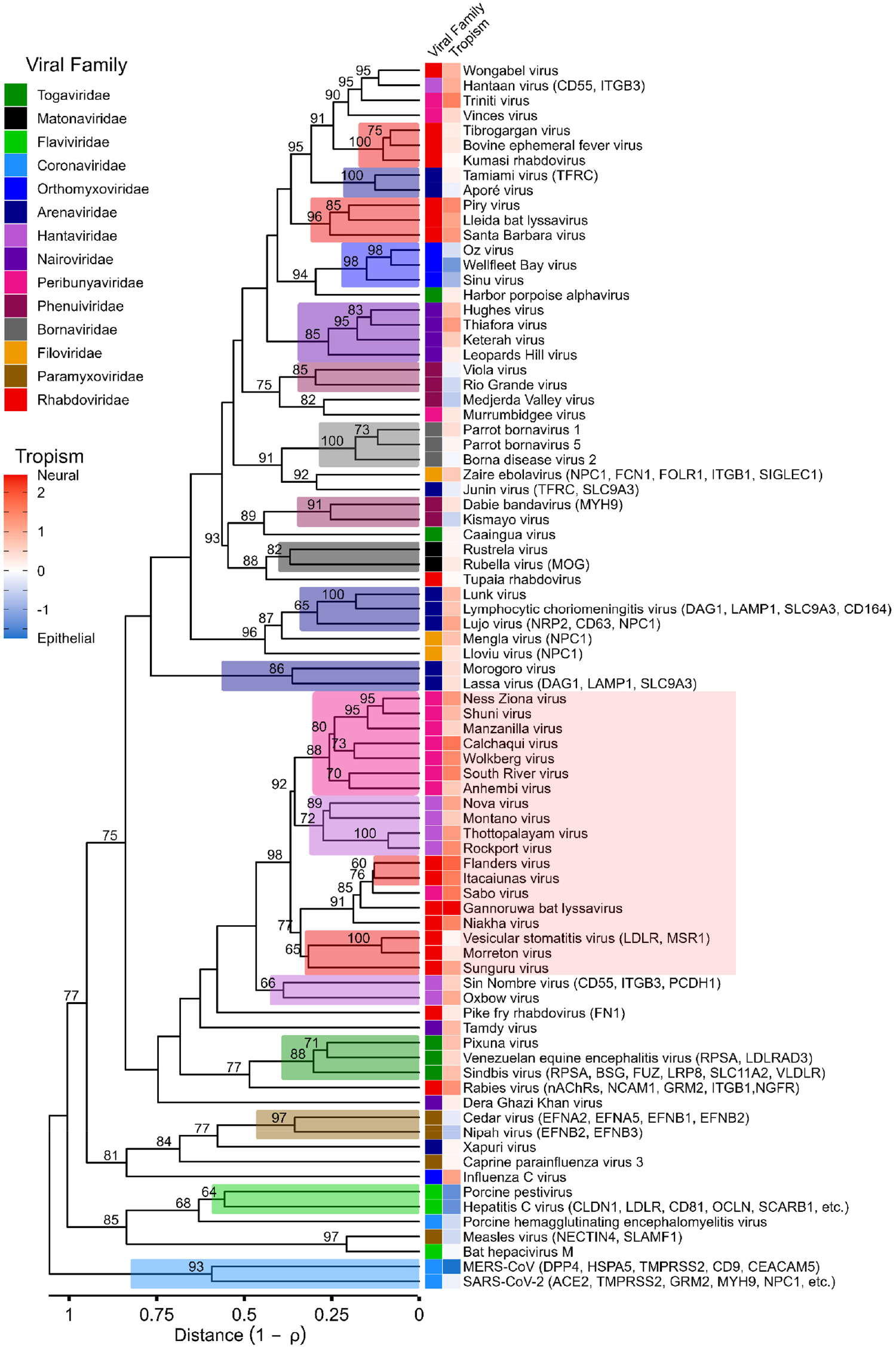
Hierarchical cluster analysis of pseudotype infectivity profiles across cell lines. The dendrogram of the 82 pseudotypes infecting at least one cell line was built using WPGMA on the Pearson correlation distance matrix derived from the infectivity correlation matrix between pairs of pseudotypes shown in **Figure S18**. Shades above branches indicate clusters of pseudotypes formed exclusively by members of the same viral family. Numbers in nodes indicate approximately unbiased bootstrap values >50%. The viral family of each virus and its tropism towards epithelial (blue) versus neuroectoderm-derived (red) cells (also shown in **Figure S19**) are displayed. The names of known viral receptors are shown in parentheses (see **Figure 4** for a complete list). The red rectangle on the right highlights a cluster of RBPs showing a significantly higher preference towards neuroectoderm-derived cells (Wilcoxon rank-sum test: P < 0.001).

### Influence of known receptors on human cellular tropism

We observed similar tropisms among some RBPs known to share cellular receptors, such as those of Nipah and Cedar viruses. In other cases, though, this similarity was weaker. For instance, Lassa virus and lymphocytic choriomeningitis virus (LCMV) are two Old-World arenaviruses that use Dystroglycan-1 (DAG1) as a specific receptor for entry^31^, but the infectivity profile of the LCMV pseudotype resembled more that of Lujo arenavirus, which uses NRP2 and CD63 instead^32^ (**Figure S20**).

We used multiple linear regression (MLR) to analyze how infectivity depended on the expression level of 65 specific receptors quantified by RNA-seq in all cell lines^33^. This included primary receptors, but also other cell surface proteins known to establish specific protein-protein interactions with RBPs. In some cases, receptor mRNA levels satisfactorily explained viral entry, such as ACE2 for SARS-CoV-2, NECTIN4 for measles, DPP4 for MERS-CoV, and EFNB2 for Nipah virus pseudotypes (MLR: P < 0.001 in all cases; **Figure 4A**; **Figure S21**). For viruses with several known receptors, we could identify those more strongly associated with tropism (**Figure 4A**). Specifically, the infection profile of the Lujo virus pseudotype was better explained by the mRNA levels of the NRP2 receptor (MLR: P < 0.001) than those of the downstream entry factor CD63. Also, infection with the Cedar virus pseudotype was positively associated with the expression of EFNB1 and EFNB2 (MLR: P < 0.001), but not of EFNA2 or EFNA5, although all have been described as receptors^34^.

**Figure 4.**
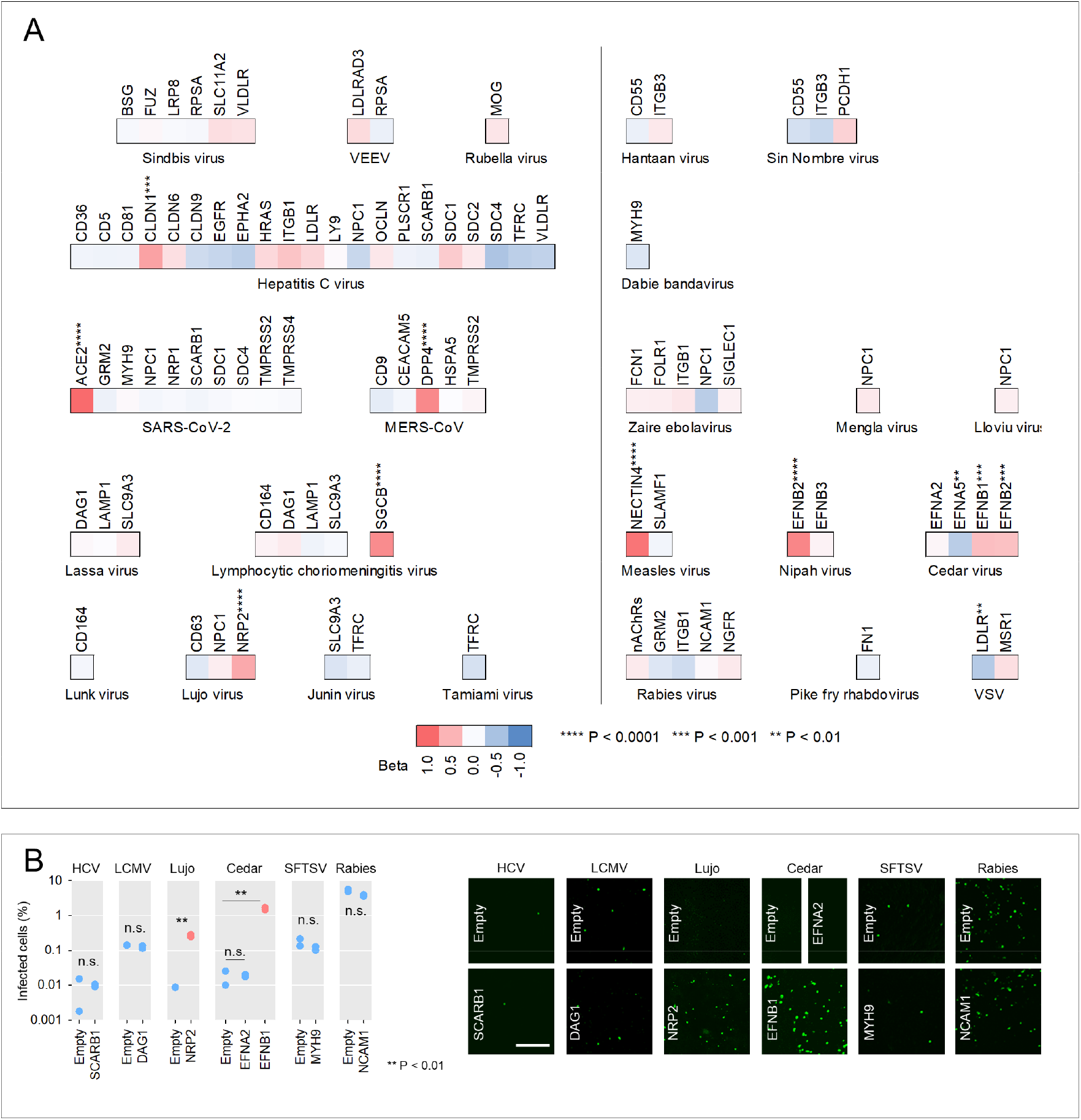
Contribution of the expression level of known receptors to RBP cellular tropism. **A**. Multiple regression analysis of pseudotype infectivity as a function of the RNA-seq expression levels of known viral receptors. For each virus, colors indicate the standardized regression coefficient. Significant coefficients are indicated with asterisks. Lectins and proteins that directly or indirectly attach to lipids were not included because we focused on proteins interacting with RBPs. For the LCMV pseudotype, the SCGB gene is included because it encodes a protein belonging to the same complex as DAG1. For the rabies virus pseudotype, the rpkm values of all genes encoding cholinergic receptor nicotinic (nAChRs) Alpha or Beta subunits were summed. Scatter plots for several significant cases are shown in **Figure S21. B**. Effect of receptor overexpression on viral entry. Cell lines were transfected with a receptor-encoding plasmid or an empty control and then infected with pseudotypes. The most appropriate cell line was chosen for each pseudotype-receptor pair based on the criteria detailed in the Methods section. These were SK-OV-3 for HCV-SCARB1, MCF7 for LCMV-DAG1 and Lujo-NRP2, ACHN for Cedar-EFNA2 and Cedar-EFNB1, UACC-62 for SFTSV-MYH9, and EKVX for Rabies-NCAM1. The percentage of infected at 24 h post-infection is shown. Two technical replicates (n = 2) were carried out for each assay. Data points shown in blue and red correspond to receptors with non-significant and significant effects on pseudotype infectivity according to the multiple linear regression shown above, respectively. Experimental results were compared to the empty controls using t-tests. Representative images are shown on the right. Scale bar: 400 µm.

However, for many other RBP-receptor combinations, the association between receptor expression levels and pseudotype infectivity was weak or absent, indicating that additional entry factors need to be considered. For instance, entry of the LCMV pseudotype did not correlate with the mRNA levels of DAG1, but did so with the expression of Sarcoglycan Beta (SGCB; **Figure S22**), another component of the dystroglycan complex. If a functional dystroglycan complex is required for LCMV entry, SCGB may effectively function as a determinant of viral tropism.

Since the frequent lack of association between receptor expression levels and viral entry was unexpected, we set out to verify this result in two ways. First, we used two quantitative proteomic datasets that included 28 and 17 of the above receptors^35,36^. In all these cases, the proteomic data confirmed the negative results obtained with RNA-seq data (**Figure S23**). Second, we transfected expression plasmids encoding different receptors and quantified the resulting changes in pseudotype infectivity (**Figure S1B**). This included two receptor-RBP pairs for which mRNA levels were significantly associated with infection in the above MLR analyses (Cedar-EFNB1 and Lujo-NRP2), versus five with no apparent association (HCV-SCARB1, LCMV-DAG1, Cedar-EFNA2, SFTSV-MYH9, and Rabies-NCAM1; **Figure 4B**). Confirming the results obtained with omics data, only EFNB1 and NRP2 overexpression increased infection by Cedar and Lujo pseudotypes, respectively. This confirms that, in many cases, viral tropism is not restricted by lack of receptor expression.

### Role of broad-range entry determinants

Most virus-specific receptors are currently unknown, which hampers the study of the molecular determinants of viral entry in less-studied viruses. We could nevertheless investigate the effects of 27 cellular proteins known to broadly influence viral entry, including caveolins (CAVs^37^), clathrins (CLTs^38^), dynamins (DNMs^39^), vimentin (VIM^40^), tetraspanins (TSPANs^41^), interferon-induced transmembrane proteins (IFITMs^42^), serine incorporators (SERINCs^43^), lymphocyte antigen 6 family member E (LY6E^44^), nuclear receptor coactivator 7 (NCOA7^45^), cholesterol-25-hydroxylase^46^, and centaurin-alpha 2 protein^47^. Viral entry dependence on each of these factors was explored using Pearson correlation coefficients between pseudotype infectivity and host gene mRNA levels across cell lines (**Figure 5A**). The strongest signal corresponded to IFITMs, which function as broad-range antiviral proteins. Indeed, we found that 43 of the 82 pseudotypes showed a significantly negative correlation (P < 0.01) between infectivity and the expression of at least one of these three genes. However, this pattern was strongly dependent on the viral family (one-way ANOVAs: P < 0.001). The most negative associations corresponded to peribunyaviruses, whereas the tropism of coronavirus RBPs was not of poorly dependent on IFITM levels (**Figure 5B**).

**Figure 5.**
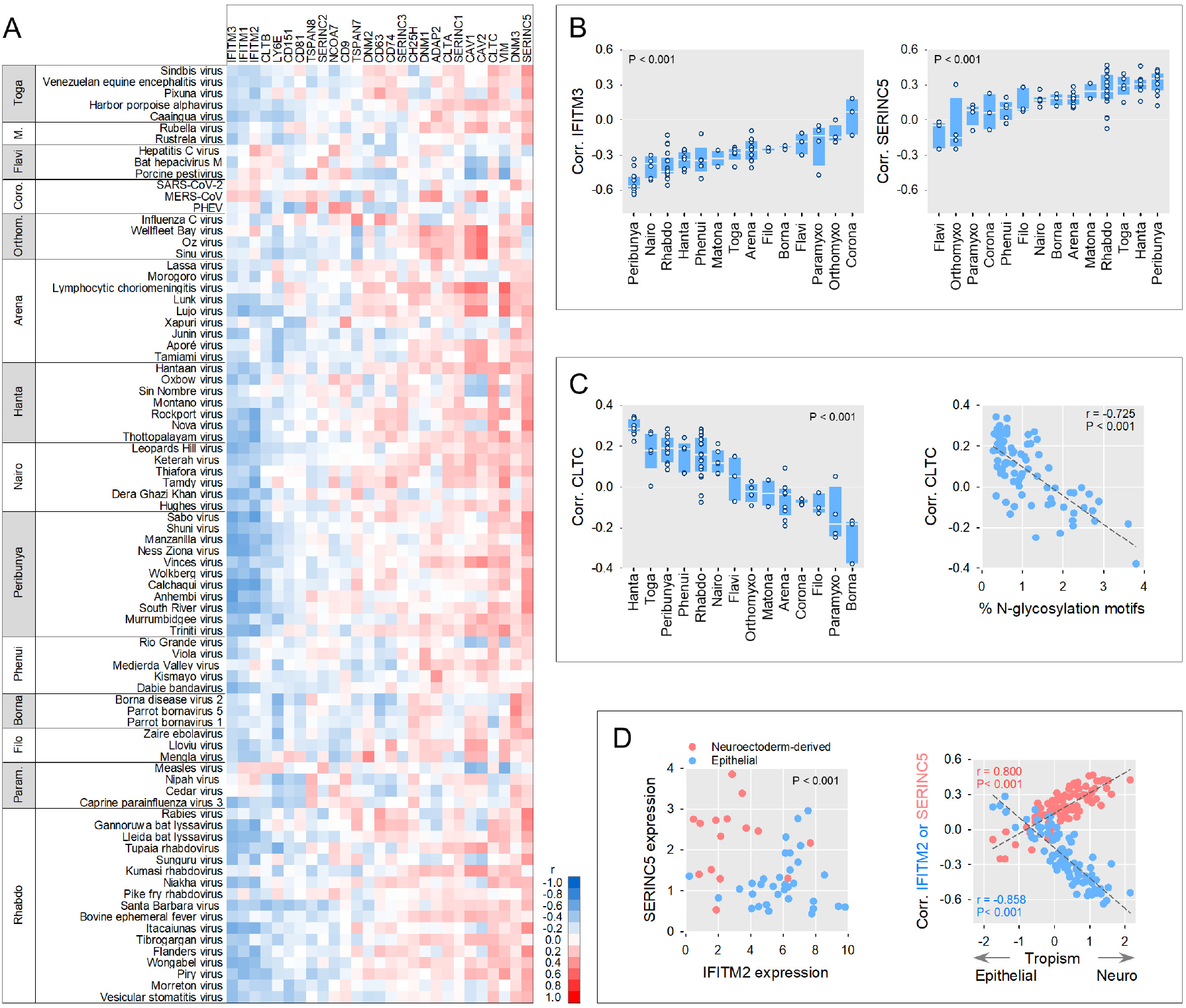
Broad-range determinants of human cell entry across animal viruses. **A**. For 27 host factors known to influence viral entry and the 82 pseudotypes, Pearson correlations between pseudotype infectivity and gene expression level across cell lines are shown. Infectivity and host gene expression levels were measured as log_2_(R+1) and log_2_(rpkm+1), respectively. The heat map indicates positive (red) or negative (blue) correlations, as shown in the scale bar. **B**. Across-family variation in these correlations for IFITM3 (left) and SERINC5 (right), the two host factors showing the strongest negative and positive associations with pseudotype infectivity across viruses, respectively. Boxes show the median (white line), 25th and 75th percentiles, and dots show data points for individual viruses. Families are sorted according to their mean correlation values, in ascending order. The significance of ANOVA tests for differences among families is shown. **C**. Left: across-family variation in the correlation between CLTC expression and pseudotype infectivity. Families are sorted according to their mean correlation values in descending order. The significance of an ANOVA test for differences among families is shown. Right: correlation between this correlation and the percentage of N-glycosylation sequence motifs in viral RBPs predicted using NetNGlyc (https://services.healthtech.dtu.dk/services/NetNGlyc-1.0/) (shown in **Table S1**). Pearson r coefficients (n = 47) and p-values are indicated **D**. Left: SERINC5 and IFITM2 expression levels, measured as log2(rpkm+1) in epithelial (blue) versus neuroectoderm-derived (red) cells. Differential expression was significant for both genes (Mann-Whitney tests: P < 0.001). Right: Correlation between SERINC5 (red) or IFITM2 (blue) expression and pseudotype infectivity, shown against the RBP preference for epithelial or neuroectoderm-derived cells. This preference was calculated as the difference between the average log_2_(R+1) values in the two cell types. Higher values indicate a preference for neuroectoderm-derived cells. Since neuroectoderm-derived cells tend to express higher levels of SERINC5 and lower levels of IFITM2, RBPs more strongly restricted by IFITM2 and favored by SERINC5 tended to be more neurotropic. Pearson r coefficients and p-values are indicated (n = 47).

In contrast to IFITMs, positive correlations with pseudotype infectivity were found for several endocytosis factors known to be exploited by viruses, including CAV1 and CAV2, the clathrin heavy chain (CTLC), and DNM3. However, CLTC dependence was conditional to the density of N-glycosylation sequence motifs present in the RBPs (r = -0.725, P < 0.001, **Figure 5C**). Recent work has shown that glycosylation inhibits the endocytosis of cellular transmembrane proteins^48^, and our data reveal that this may also apply to viral clathrin-mediated viral endocytosis. Predicted RBP glycosylation as well as the association between CLTC expression and pseudotype infectivity also varied significantly across viral families (one-way ANOVA: P < 0.001; **Figure 5C**).

We found unexpected associations between the expression levels of some host genes and viral entry. First, LY6E has been shown to enhance viral infection, mainly in flaviviruses, but also in some togaviruses, orthomyxoviruses, and retroviruses^49^. In contrast, we found that LY6E mRNA levels correlated negatively with pseudotype infectivity for most viral families, the exception being flaviviruses. Our observations are consistent with work suggesting an antiviral role for this gene^44^. Second, SERINC5, which has been identified as a factor blocking the entry of HIV-1, other retroviruses, and Influenza A virus^43,50^, surprisingly showed an overall positive association with viral entry, again with ample variation across families (one-way ANOVA: P < 0.001; **Figure 5B**). Moreover, we found that viruses showing a stronger association between SERINC5 levels and RBP-mediated entry were those exhibiting a more marked RBP preference for neuroectoderm-derived cells (**Figure 5D**). Analysis of additional candidate host genes associated with this tropism led to the identification of IFITM2. Both genes were differentially expressed in epithelial versus neuroectoderm-derived cells (**Figure 5D**), suggesting that constitutive levels of SERINC5 and IFITM2 contribute to determining the cellular tropism of animal viruses.

## Discussion

Our results challenge the view that incompatibilities between viral entry factors across host species constitute a major barrier to zoonosis, since most RBPs of enveloped RNA viruses mediated entry into human cells. This implies that entry factors are sufficiently conserved to permit cross-species viral transmission. Coronaviruses may constitute a remarkable exception since most their RBPs failed to mediate entry into any of the human cell lines tested or showed a narrow cell tropism. Known coronavirus receptors such as ACE2 and DPP4 were expressed in few cell lines, in contrast to ubiquitous viral receptors such as DAG1, LDLR, NPC1, and EFNA1, among others. Moreover, coronavirus entry involves additional RBP-specific factors such as host proteases, which are required for spike activation and play a key role in unlocking the human infectivity of some coronaviruses^14,51,52^.

Our results also reveal that the presence of a receptor is not a limiting factor for viral entry in most cases, and that other cellular factors can often effectively determine viral tropism. Some receptors may be necessary for viral entry but be expressed ubiquitously whereas, in other cases, entry may be achieved through multiple alternative receptors and pathways. Besides, viral infectivity can also be strongly influenced by broadly acting host factors, and we show that these may play a role in determining intra-host viral tropism. IFITM expression exhibited a systematic negative association with RBP-mediated entry, particularly for peribunyaviruses. Although IFITMs are well-characterized broad-spectrum antiviral factors, how their effects vary across viral families remains poorly understood. RBPs may show different levels of susceptibility to IFITM-mediated restriction or, alternatively, the observed variation across families may reflect differences in the contribution of IFITMs to RBP tropism relative to other entry factors, such as virus-specific receptors. We also found viral family-dependent effects for other broad-range entry factors such as SERINC5 and the Cathrin heavy chain, and identified a link between RBP glycosylation and clathrin-mediated viral endocytosis that should be further explored in future work.

The in-depth characterization of entry factors may allow the development of broad-spectrum host-targeted antiviral drugs and improve our preparedness against novel emerging viruses. Indeed, a battery of antivirals active against viruses from different families would allow bypassing the development of virus-specific drugs and respond more quickly to emerging viruses^53^. Many research teams attempt to predict which viral species are more likely to emerge in humans^54^, and several studies have identified viral features associated with cross-species transmission and zoonosis^6–9^. For instance, it has been shown that machine learning applied to viral genomic data can predict whether a virus is vector-borne^55^ or is at risk of infecting humans^56^. However, because functional information on wildlife viruses is often lacking, data on their ability to infect human cells are rarely included in such predictions. Our study contributes to filling this information gap by characterizing the human cellular tropism of a large number of wildlife viruses.

Tumor cells are widely used models in virology, including some NCI-60 cells such as A549. Although these cells show abnormal cell physiology, they allow testing whether animal viruses are compatible with the human version of native-host entry factors. Performing such a systematic screening of human infectivity in human primary cells would be technically much more complex. A major advantage of the NCI-60 panel is that their gene expression profile has been extensively characterized, which allowed us to explore the molecular determinants of viral entry. Moreover, we found that the expression levels of cell surface genes in these cells correlated well with those reported for normal tissues. Thus, the NCI-60 panel offers an experimentally tractable system for high-throughput cell culture screening and infection, and a valuable tool for investigating virus-host interactions, as shown previously^57,58^.

Here, we have focused on viral entry due to its presumed importance in zoonosis and because pseudotyping allows the study of entry mechanisms even for viruses that have never been isolated, as is the case for most of the viral species included in our analysis. However, entry is only the first step that a virus must complete to productively infect a cell, produce progeny virions, and spread at the intra-host and population levels. These other aspects of infection should also be studied to achieve a more comprehensive view of zoonotic risks. Indeed, our results suggest that the post-entry stages of the infection cycle, as well as epidemiological and ecological factors, may be more critical determinants of viral zoonosis than viral entry.

## Methods

### RBP sequence retrieval and phylogenetic analyses

For each viral family, viral species were retrieved from both the ICTV (https://ictv.global) and the NCBI RefSeq (https://www.ncbi.nlm.nih.gov/refseq/) databases and one representative of each species was selected. In some cases (e.g., Betacoronavirus-1), more than one representative was included due to special relevance or different origins. The genomes of the selected virus were retrieved from the NCBI database, their CDSs translated into protein and annotated with InterProScan (https://www.ebi.ac.uk/interpro/) to identify the protein of interest. In case of the flavivirus and nairovirus polyproteins, the region of interest was retrieved directly from NCBI Entrez. Different protein alignment methods were compared and Clustal Omega (ebi.ac.uk/jdispatcher/msa/clustalo) was chosen. The best amino acid substitution model was then selected using ProtTest3 (github.com/ddarriba/prottest3) and used for building a maximum-likelihood tree was using RaxML (github.com/stamatak/standard-RAxML).

### Synthesis of RBP sequences

For each viral family, viral species of interest were selected across different clades to cover as much viral diversity as possible. The full RBP sequence was synthesized, with the exception of Flaviviridae and Togaviridae where only parts of the structural polyprotein gene were synthesized (C-terminal part of C-E1E2 and E3E26kE1, respectively). All sequences were codon-optimized for human expression, except for VSV, Ebola and Tamiami virus. RBP-coding DNA was cloned into a pcDNA3.1-C-HisTag vector (Genscript). For paramyxoviruses, fusion proteins were cloned into a pcDNA3.1-C-Flag vector, and attachment glycoproteins were cloned into a pcDNA3.1-N-HisTag vector. For Nipah virus, the F and G proteins were cloned in a single pCI-Neo-G-IRES-F plasmid. The Ebola GP-encoding plasmid was obtained from Addgene. Information about RBPs is provided in **Table S1**.

### Pseudotyping

T75 flasks were coated with poly-D-lysine (Gibco) for 2 h at 37ºC, washed with water, and seeded with 8 x 10^6^ HEK293T cells. The following day, cells were transfected with 30 µg of viral glycoprotein expression plasmid using Lipofectamine 2000 (Invitrogen) following the manufacturer’s instructions. In the case of paramyxoviruses, cells were transfected with a mixture of 15 µg of fusion protein- and 15 µg of hemagglutinin/glycoprotein-expressing plasmids. To produce bald pseudotypes to be used as negative controls in infection experiments, cells were transfected with an empty pcDNA3.1 vector. At 24 h post-transfection, cells were inoculated at a multiplicity of infection (MOI) of 3 infectious units per cell for 1 h at 37ºC with a VSV encoding GFP, lacking the glycoprotein gene G (VSVΔG-GFP), and previously pseudotyped with G. Cells were washed three times with PBS and 8 mL of DMEM supplemented with 2% FBS were added. Supernatants containing pseudotypes were harvested 24 h later, cleared by centrifugation at 2000 g for 10 min, passed through a 0.45 µm filter, aliquoted, and stored at -80C.

### Western blotting

A 1 mL volume of supernatant containing pseudotype was pelleted by centrifugation at 30,000 g for 2 h at 4ºC and lysed in 30 µL of NP-40 lysis buffer (Invitrogen) for 30 min on ice. Approximately 10^6^ pseudotype-producing cells were lysed for 30 min on ice in 100 µL of NP-40 lysis buffer. Lysates were cleared by centrifugation at 15,000 g for 10 min at 4ºC. Cleared lysates were mixed with 4X Laemlli buffer (Bio-Rad) supplemented with 10% β-mercaptoethanol and denatured at 95ºC for 5 min. Proteins were separated by SDS-PAGE on a 4-20% Mini-PROTEAN TGX Gel (Bio-Rad) and transferred onto a 0.45 µm PVDF membrane (Thermo Scientific). Membranes were blocked for 1 h at room temperature in TBS-T (20 mM tris, 150 mM NaCl, 0.1% Tween-20, pH 7.5) supplemented with 3% Bovine Serum Albumin (BSA; Sigma). Membranes were then incubated for 1 h at room temperature with the following primary antibodies: mouse anti-His-Tag (dilution 1:1,000, clone HIS.H8, Invitrogen MA121315), mouse anti-Flag (dilution 1:1000, clone M2, Sigma-Aldrich F1804), mouse anti-VSV-M (dilution 1:1000, clone 23H12, Kerafast EB0011), or rabbit anti-GAPDH (dilution 1:3000, Sigma-Aldrich ABS16). Membranes were washed 3 times with TBS-T and incubated for 1 h at room temperature with an HRP-conjugated anti-mouse (dilution 1:50,000, Invitrogen, G-21040) or anti-rabbit (dilution 1:50,000, Invitrogen, G-21234) secondary antibody. After 3 washes in TBS-T, the signal was revealed with SuperSignal West Pico PLUS (Thermo Scientific) following the manufacturer’s instructions. Images were acquired on an ImageQuant LAS 500 (GE Healthcare) and analyzed with Fiji software.

### Selection of NCI-60 cell lines

The NCI-60 panel (dtp.cancer.gov/discovery_development/nci-60) was obtained from the National Cancer Institute. The panel consists of 60 well-characterized cancer cell lines from various origins. Only adherent cell lines were included to facilitate imaging, thus excluding the six leukemia cell lines (MOLT-4, RPMI-8226, SR, CCRF-CEM, HL-60, K562), one lung cell line (NCI-H322M) and one colon cell line (COLO 205). The HCC2998 cell line was also excluded due to poor cell growth. For convenience for high-throughput analyses, the number of cell lines was reduced to 48 by excluding three other cell lines (HS 578T, LOX IMVI, KM12). Information about the 48 cell lines is provided in **Table S2**.

### Verification of the identity of each NCI-60 cell line

The 48 NCI-60 cell lines included in our analyses were authenticated by Short Tandem Repeat (STR) genotyping. Briefly, genomic DNA was extracted using the PureLink Genomic DNA Mini Kit (Invitrogen) following the manufacturer’s instructions. Genomic DNA concentration was quantified using a NanoDrop One spectrophotometer (Thermo) and 1 ng of DNA was amplified by PCR using the AmpFℓSTR Identifiler Plus PCR Amplification Kit (Applied Biosystems) following the manufacturer’s instructions. Amplified fragments were analyzed by capillary electrophoresis using a 3730 XL DNA Analyzer (Applied Biosystems). Chromatograms were analyzed using the Osiris software (ncbi.nlm.nih.gov/osiris). Results were compared to the STR profiles of the NCI-60 cell lines available online (web.expasy.org/cellosaurus) using a relaxed similarity metric. All cell lines were correctly identified.

### NCI-60 cell culture

Cells were cultured in 12-well plates in RPMI (Gibco) supplemented with 10% FBS (Gibco), 10 units/mL penicillin, 10 µg/mL streptomycin (Gibco), 250 ng/mL amphotericin B (Gibco) and 5 µg/mL prophylactic plasmocin (InvivoGen). Before passaging, plates were imaged in the Incucyte SX5 Live-Cell Analysis System (Sartorius). Cell confluence was quantified automatically with the Incucyte Analysis software using the Artificial Intelligence Confluence segmentation algorithm with a cleanup hole fill parameter of 100 µm^2^ and filtering out segments smaller than 200 µm^2^. Cells were washed with PBS, detached with trypsin (Gibco), complete culture medium was added, and cells were dispensed in new 12-well plates for maintenance and in 96-well plates at a 30% confluence for infection experiments the following day. Passaging was performed every two or three days. Cell lines were regularly shown to be free of mycoplasma contamination by PCR.

### Infections

Pseudotypes were mixed 1:1 with an anti-VSV-G monoclonal antibody to remove residual VSV-G and incubated for 20 min at 37ºC. Cell culture medium was removed and cells were inoculated with 50 µL of the antibody-treated pseudotypes. Plates were incubated for 2 h at 37ºC and 50 µL of RPMI supplemented with 5% FBS were added to each well. After 18-24 h, cells were imaged in the Incucyte SX5 Live-Cell Analysis System (Sartorius). Cell confluence and the percentage of GFP-positive area were quantified automatically with the Incucyte Analysis software. Each experimental block consisted of four 96-well plates containing the 48 cell lines arranged in 12 × 4 columns. Six rows of each plate were used to test different pseudotypes, and two rows were used to carry out two internal controls. First, a blank row inoculated with a bald pseudotype was used to measure the background signal resulting from cell auto-fluorescence or residual VSV-G-pseudotyped particles. The values obtained in these negative controls were subtracted from the corresponding measurements. Second, a row inoculated with VSVΔG-GFP pseudotyped with G, which allowed us to assess inter-assay reproducibility for a given RBP. Each of the 102 pseudotypes was assayed twice in experimental blocks performed on different days.

### Quantitation of pseudotype infectivity

The proportion of infected cells, Q, was measured as the ratio between the GFP area and cell confluence. To correct for saturation effects in highly infected wells, we transformed Q-values into MOIs as follows: MOI = –ln (1-Q). For Q-values >0.95, an MOI of 3 infectious units per cell was assumed. To account for differences in pseudotype titer, for each pseudotype and assay, relative MOI values were calculated as R = 100 × MOI/max(MOI), i.e. as a percentage of the maximal MOI observed among the 48 cell lines. Finally, values were log-transformed as log_2_ (R+1), and the two log_2_ (R+1) data obtained for each RBP-cell combination were averaged. For the VSV-G internal controls, the median Pearson correlation coefficient between log_2_ (R+1) values from different experimental blocks was r = 0.834 and 91.2% of the individual data points were within twofold of the inter-assay average, validating the reproducibility of the assays. In addition to quantifying infection, for each of the 102 × 48 RBP-cell combinations, we obtained a dichotomous variable indicating the presence or absence of infection. The following conditions had to be met in both replicates of each RBP-cell combination for this variable to be positive: (i) R > 1; (ii) Q at least 5 times higher than in the corresponding empty-plasmid control; (iii) Q > 0.05%. Visual inspection of multiple wells was used to establish these criteria. Average log_2_ (R+1) values and positivity data are provided in **Table S3**.

### Human gene cloning

For each gene of interest, the sequence of the main human transcript (flagged as MANE Select) was retrieved from Ensembl (ensembl.org) and the NCBI RefSeq databases (ncbi.nlm.nih.gov/refseq). RNA was extracted from HEK293T cells or NCI-60 cell lines expressing the gene of interest according to RNA-seq data, using RNAzol (Sigma-Aldrich) following the manufacturer’s instructions. RNA was reverse-transcribed into cDNA using Oligo dT and SuperScript IV Reverse Transcriptase (Invitrogen) following the manufacturer’s instructions. The cDNAs were used as templates for PCR amplification using primers detailed **in Table S4**. PCR-amplified transcripts were cloned into a pcDNA3.1-C-Flag or pcDNA3.1-N-HA vector with restriction enzymes or through HiFi assembly. For restriction enzyme cloning, restriction sites were added in amplification primers. The vector and PCR products were digested with restriction enzymes and band-purified (vector) or cleaned (PCR products) using the Zymoclean Gel DNA Recovery Kit (Zymo Research) or the DNA Clean & Concentrator-5 kit (Zymo Research), respectively. Purified PCR products and the vector were mixed at a 1:3 molar ratio and ligated using T4 DNA ligase (Thermo Scientific). For HiFi cloning, the pcDNA3.1-C-Flag vector was linearized by PCR (Forward primer: 5’-GATTACAAGGATGACGACGATAAGTG-3’; Reverse primer: 5’-GGTGGCAAGCTTAAGTTTAAACGCTAG-3’). Amplification primers contained a 20-nucleotide tail overlapping with the 5’ or 3’ ends of the linearized pcDNA3.1-C-Flag vector. The linearized vector and PCR-amplified sequences were mixed at a 1:2 molar ratio and assembled using the NEBuilder HiFi DNA Assembly Master Mix (New England Biolabs) following the manufacturer’s instructions. Phusion Hot Start II High-Fidelity DNA polymerase (Thermo Scientific) was used for all PCR steps. Assembled products were transformed into NY5α competent cells (NZYtech). Correct insertion was checked by colony PCR using vector-specific primers (Forward: 5’-GAGAACCCACTGCTTACTGGC-3’; Reverse: 5’-AGGGTCAAGGAAGGCACG-3’) and the NZYTaq II 2x Green Master Mix (NZYtech). Plasmids with correct insertions were checked by Sanger (Eurofins) or whole-plasmid high-throughput sequencing (Plasmidsaurus). Additionally, correct production of the protein of interest was checked by Western Blot of HEK293T-transfected cells using an anti-Flag antibody (Sigma-Aldrich).

### Host gene overexpression assays

For each pseudotype-gene combination, the most appropriate cell line was based on the following criteria: low expression of the gene to be tested, good transfection efficiency, low infection by the virus to be tested, and good susceptibility to VSV. Cells were plated in 96-well plates. The following day, cells were transfected with the gene-encoding plasmid or an empty vector control using Lipofectamine 2000 or Lipofectamine 3000 (Invitrogen), following the manufacturer’s instructions. A control for transfection efficiency was included (GFP expression plasmid). After 20-24 h, pseudotypes were mixed 1:1 with an anti-VSV-G monoclonal antibody and incubated for 30 min at 37ºC. Cell culture medium was removed and cells were inoculated with 50 µL of antibody-treated pseudotypes. Plates were incubated for 2 h at 37ºC and 50 µL of RPMI supplemented with 5% FBS were added to each well. After 18-24 h, cells were imaged in the Incucyte SX5 Live-Cell Analysis System (Sartorius). Cell confluence and the percentage of GFP-positive area were quantified automatically with the Incucyte Analysis software. The proportion of infected cells was calculated as the ratio between the GFP area and cell confluence.

### RNA-seq and proteomics data

A processed RNA-seq dataset was downloaded from the CellMiner website (discover.nci.nih.gov/cellminer/loadDownload.do, RNA-seq - composite expression file), as well as proteomic data available for a subset of virus receptors (SWATH Mass spectrometry – Protein file). An additional proteomics was obtained from ebi.ac.uk/pride/archive/projects/PXD005940. These omics data were available for all the cell lines except MDA-MB-468. Gene expression values were available as log_2_(rpkm+1). RNA-seq data averaged over 40 human tissues were downloaded from the Human Protein Atlas (proteinatlas.org/about/download, rna_tissue_hpa file).

### Statistical analyses

Differences in the proportion of infected cell lines among viral families were tested using a logistic regression analysis, in which the dependent variable was the presence/absence of infection in each of the 102*48 RBP-cell combinations and the independent variable was the viral family. Wilcoxon rank-sum tests were used to compare the fraction of cells infected by pseudotypes of human-infective versus non-human viruses, and of arthropod-infective versus non-arthropod viruses. Wilcoxon tests were also used to compare the average log_2_(R+1) values in epithelial versus neuroectoderm-derived cells. A hierarchical cluster analysis was carried out to classify pseudotypes according to their similarly in infectivity profiles, measured as log2(R+1) across the 48 cell lines. Several distance metrics (Pearson correlation distance, cosine distance and Euclidean distance) and linkage methods (UPGMA, WPGMA, Ward and Neighbor Joining) were tested. The dendrogram obtained by Pearson correlation distance (1-ρ) and WPGMA linkage was the one that best recapitulated the viral phylogeny based on the average size and number of viruses included in groups monophyletic for the viral family. The stability of dendrogram nodes was evaluated applying multiscale bootstrap resampling upon infectivity data using pvclust R package (https://github.com/shimo-lab/pvclust). Multiple linear regression was used to estimate the relative contribution of different virus receptors to observed infectivity data, where the dependent variable was log_2_(R+1) across all cell lines and the independent variable were the standardized log_2_(rpkm+1) data for receptor. In gene overexpression experiments, mean GFP signals were compared using t-tests with log-transformed data. The association of infectivity profiles with the expression of broad-range entry factors was evaluated using Pearson correlations between log_2_(R+1) and log_2_(rpkm+1) data across all cell lines. All statistical tests were two-sided. Statistical analyses were carried out with R and SPSS v28.

## Data availability

Features of the RBPs analyzed in this study are available in **Table S1**. Features of the 48 cell lines of the NCI-60 panel are available in **Table S2** and at dtp.cancer.gov/discovery_development/nci-60/cell_list.htm. Pseudotype infection data are available in **Table S3**. Primer information is available in **Table S4**. NCI-60 omics data are available at discover.nci.nih.gov/cellminer/loadDownload.do, and at ebi.ac.uk/pride/archive/projects/PXD005940. RNA-seq data from the Human Protein Atlas are available at proteinatlas.org/about/download.

## Acknowledgments

We thank Carlos Baeza-Delgado and Raquel Martínez-Recio for technical assistance, and Jorge Moreno for helpful comments. This work was financially supported by an ERC Advanced Grant (101019724—EVADER) and a grant from the Spanish Ministerio de Ciencia e Innovación (PID2020-118602RB-I00—ZooVir) to R.S. J.D. is the recipient of an EMBO postdoctoral fellowship (ALTF 140-2021) and a Marie Skłodowska-Curie Actions Postdoctoral Fellowship (101104880).

## Authors’ contributions

J.D. and R.S. designed research; J.D., I.A.M and A.V.R. performed research; J.D., I.A.M and R.S. analyzed data; J.D. and R.S. wrote the paper; R.S. provided funding.

