## Supplementary information for "Viral entry is a weak barrier to zoonosis"

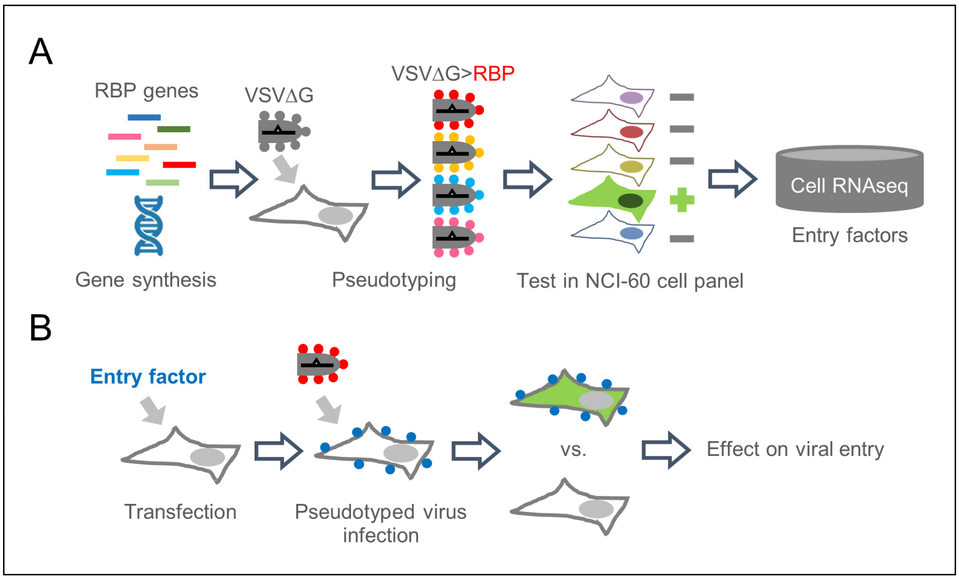


**Figure S1. Experimental workflow. A.** RBP sequences were obtained from public databases and used for gene synthesis. RBPs were used for creating VSV pseudotypes by transfection in HEK293T cells and transduction with a VSV deleted for the envelope gene G (VSVΔG-GFP) and pseudotyped with G. Pseudotypes carrying the RBPs of interest were assayed for infectivity in 48 human cell lines from the NCI-60 panel. Infectivity profiles were compared to cellular gene expression data to evaluate the contribution of different host proteins including virus-specific receptors and broad-range entry factors. **B.** The effect of some virus receptors on viral entry was tested by transfecting relevant genes and infecting with VSV pseudotypes.

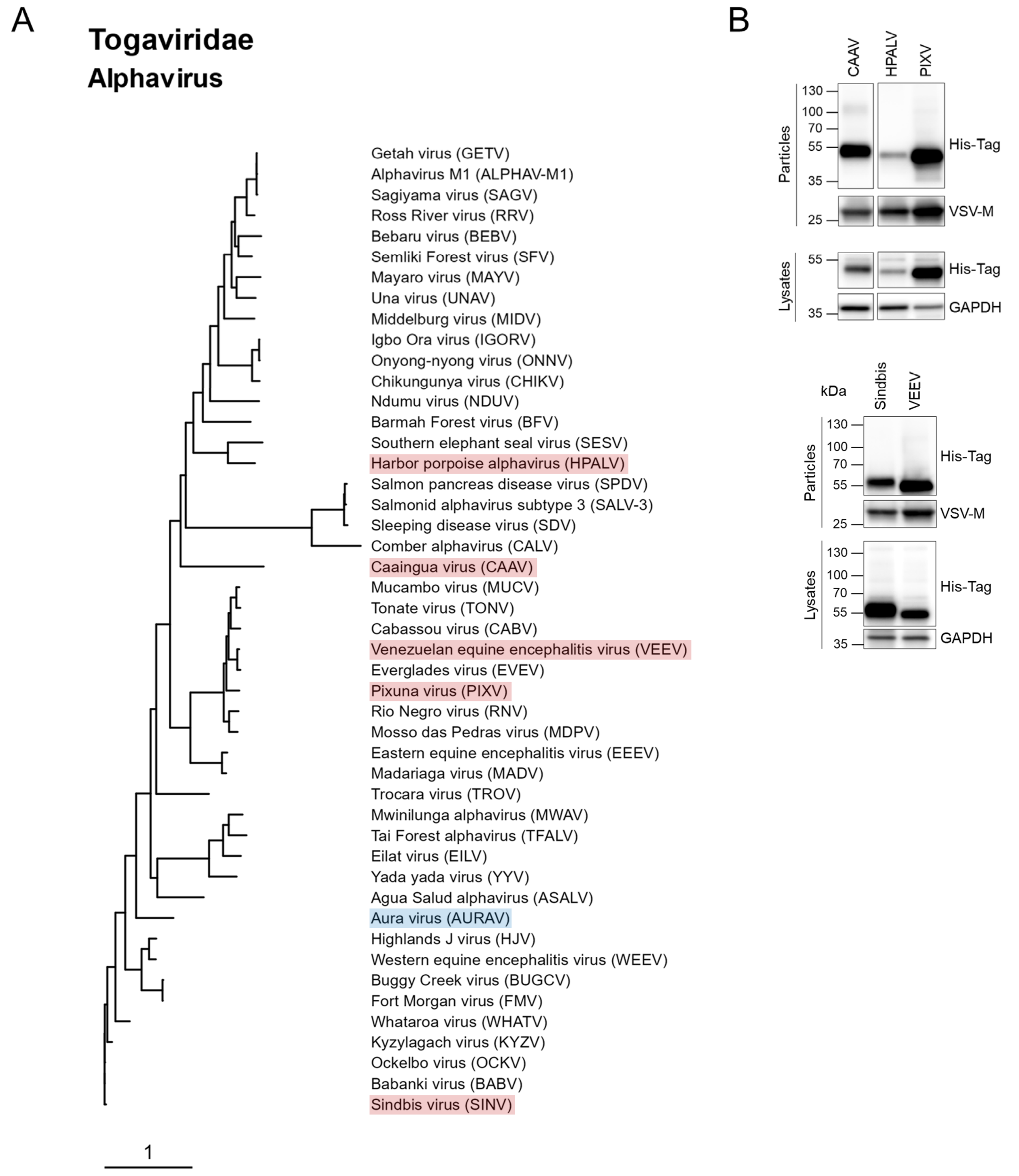


**Figure S2. Togaviridae structural protein phylogeny for the alphavirus genus and Western Blot confirmation of correct pseudotype production. A.** Maximum-likelihood phylogenetic tree of the CPE3E26kE1 polyprotein. Species included in our analysis are shaded in red. Species shaded in blue are viruses for which pseudotype production was attempted but unsuccessful. **B.** The E3, E26k and E1 proteins were expressed. A His-Tag added to the E1 protein C-terminus was used for detection in producer cells and to show incorporation into VSV pseudotypes. GAPDH and VSV-M were used as loading controls for producer cells and viral pseudotypes, respectively.

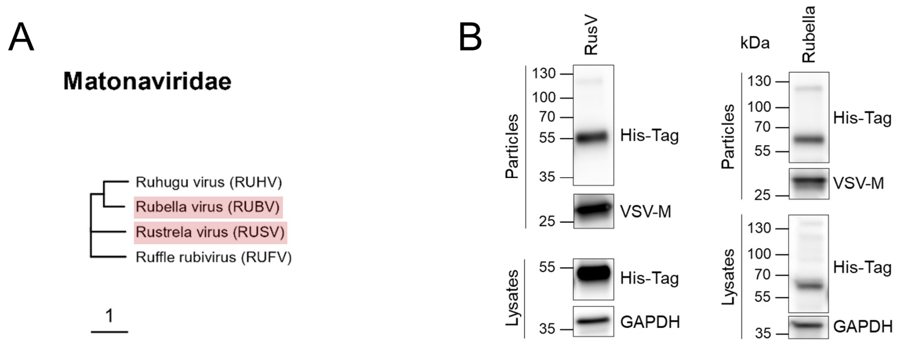


**Figure S3. Matonaviridae structural protein phylogeny and Western Blot confirmation of correct pseudotype production. A.** Maximum-likelihood phylogenetic tree of the CPE2E1 polyprotein. Species included in our analysis are shaded in red. **B.** The C, E2 and E1 proteins were expressed. A His-Tag added to the E1 C-terminus was used for detection in producer cells and to show incorporation into VSV pseudotypes. GAPDH and VSV-M were used as loading controls for producer cells and viral pseudotypes, respectively.

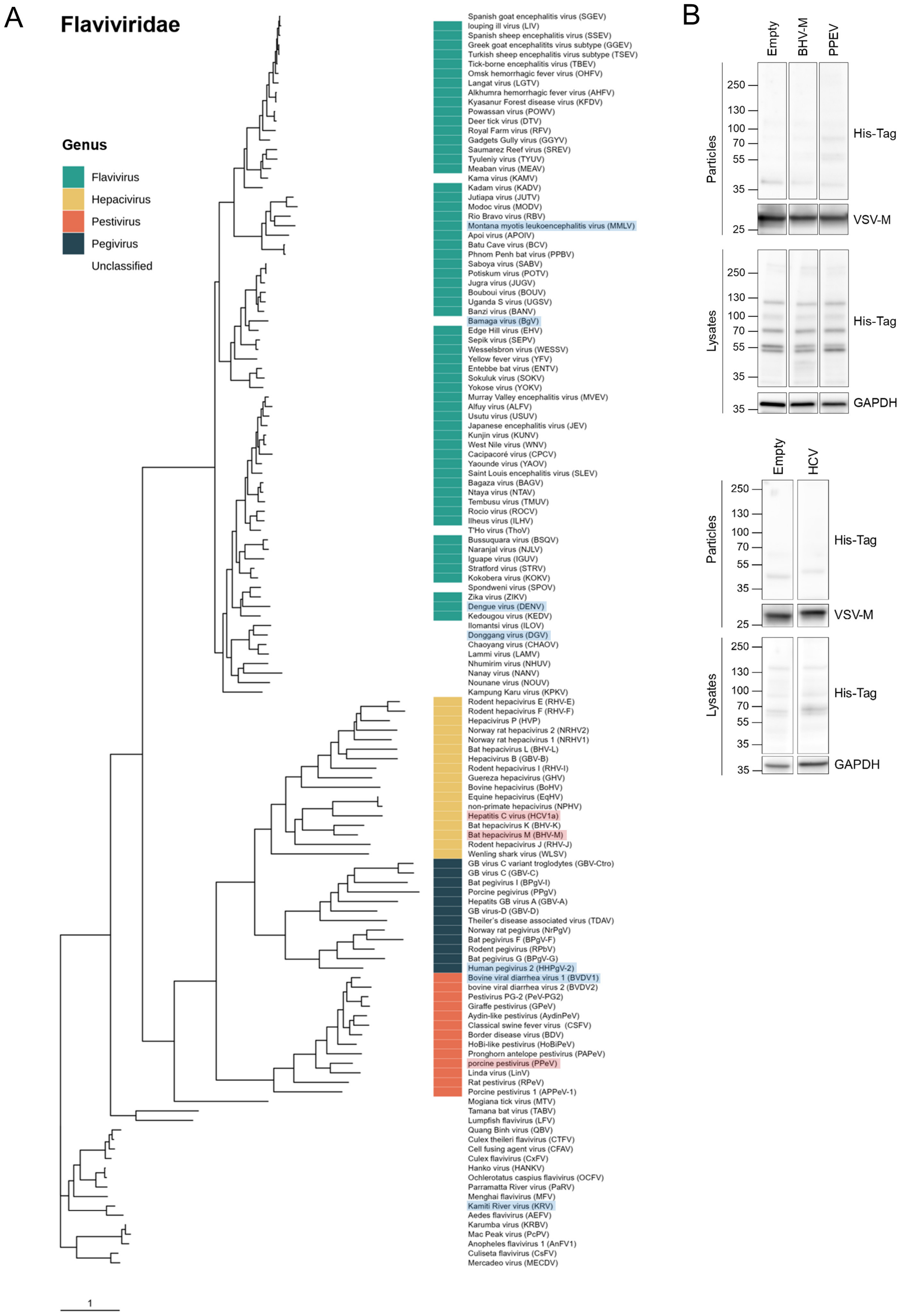


**Figure S4. Flaviviridae envelope proteins phylogeny and Western Blot confirmation of correct pseudotype production. A.** Maximum-likelihood phylogenetic tree. Species included in our analysis are shaded in red. Species shaded in blue are viruses for which pseudotype production was attempted but unsuccessful. **B.** The C terminus of the C protein and the entire E1E2 (HCV and BHV-M) and $E^{rns}$E1E2 (PPEV) proteins were expressed. A His-Tag added to the E2 protein C-terminus was used for detection in producer cells and to show incorporation into VSV pseudotypes. GAPDH and VSV-M were used as loading controls for producer cells and viral pseudotypes, respectively.

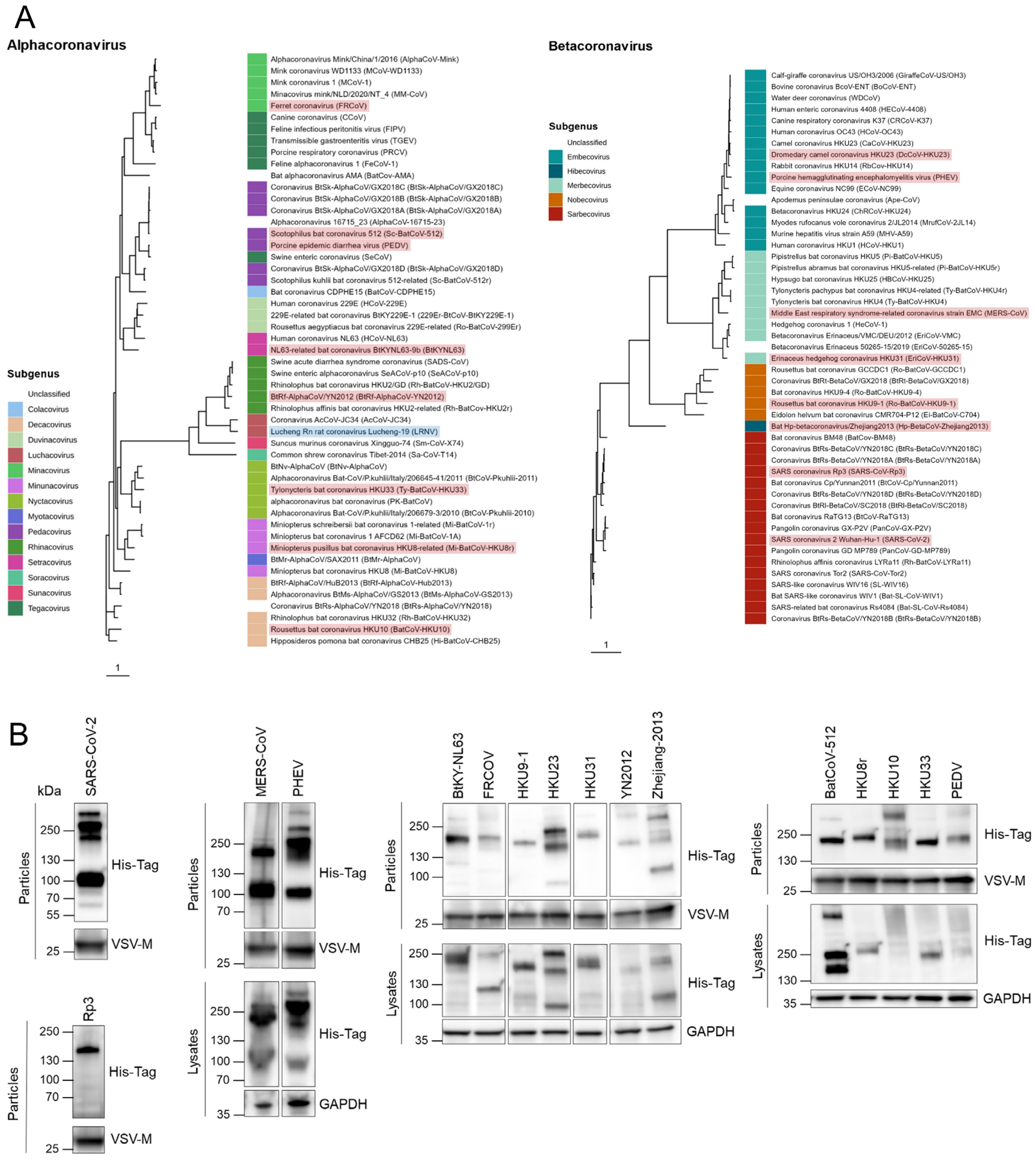


**Figure S5. Coronaviridae spike protein phylogeny and Western Blot confirmation of correct pseudotype production. A.** Separate maximum-likelihood phylogenetic trees for the alpha and betacoronaviruses. Species included in our analysis are shaded in red. Species shaded in blue are viruses for which pseudotype production was attempted but unsuccessful. **B.** A His-Tag added to the spike protein C-terminus was used for detection in producer cells and to show incorporation into VSV pseudotypes. GAPDH and VSV-M were used as loading controls for producer cells and viral pseudotypes, respectively.

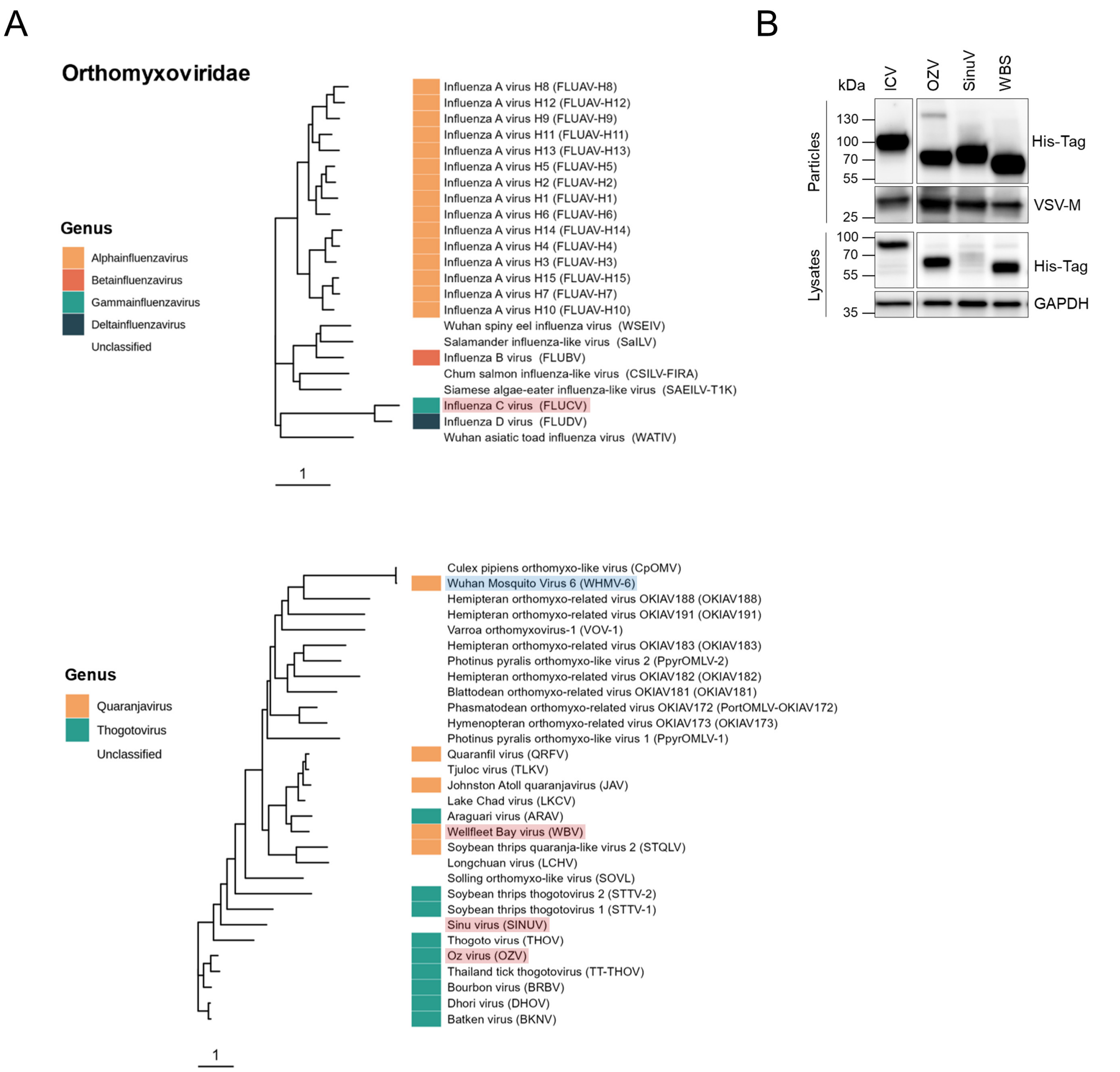


**Figure S6. Orthomyxoviridae RBP phylogeny and Western Blot confirmation of correct pseudotype production. A.** Maximum-likelihood phylogenetic tree of the HA (top tree) or GP (bottom tree) depending on genus. Species included in our analysis are shaded in red. Species shaded in blue are viruses for which pseudotype production was attempted but unsuccessful. **B.** A His-Tag added to the HA or GP C-terminus was used for detection in producer cells and to show incorporation into VSV pseudotypes. GAPDH and VSV-M were used as loading controls for producer cells and viral pseudotypes, respectively.

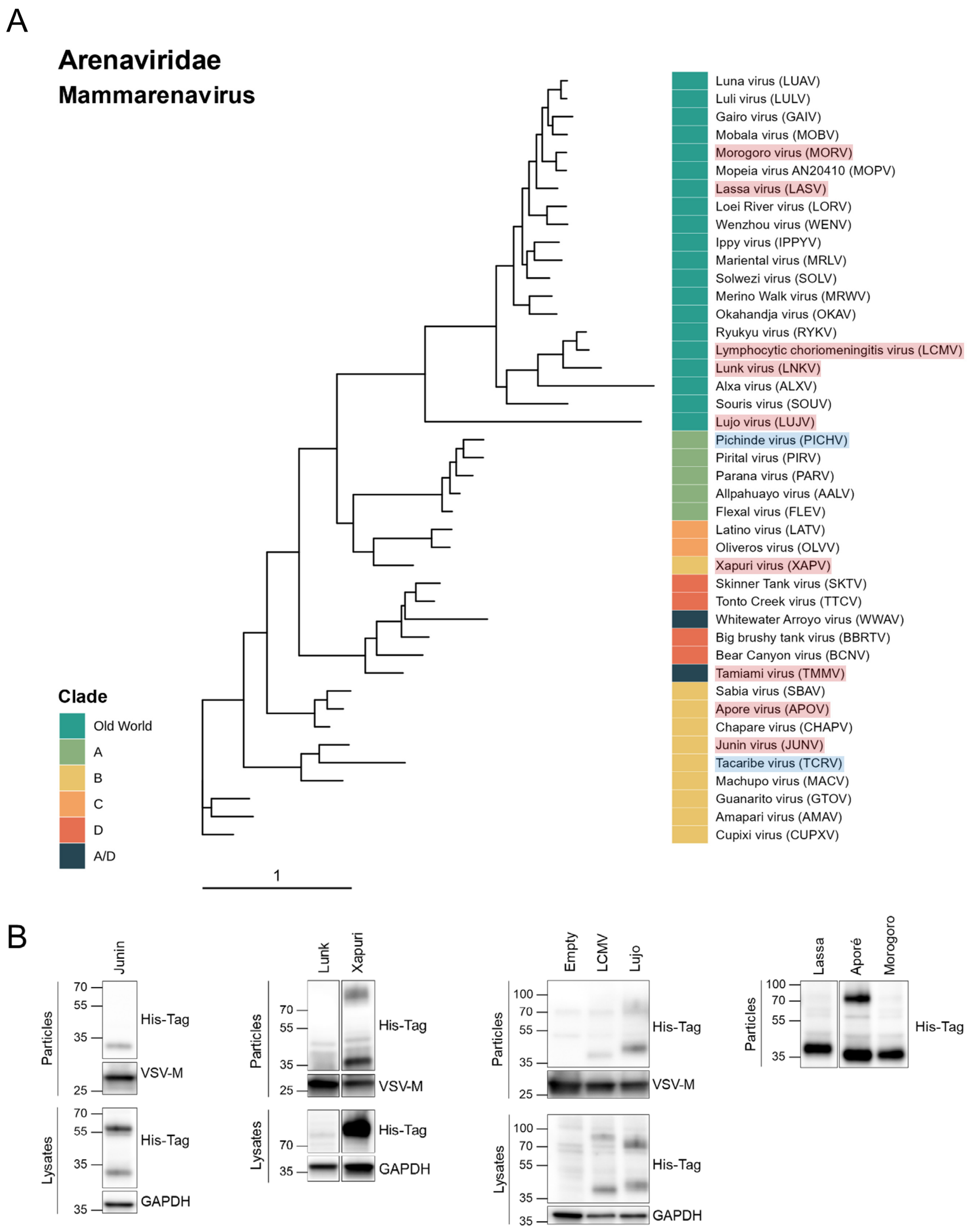


**Figure S7. Arenaviridae GP phylogeny for the mammarenavirus genus and Western Blot confirmation of correct pseudotype production. A.** Maximum-likelihood phylogenetic tree of the RBPs. Species included in our analysis are shaded in red. Species shaded in blue are viruses for which pseudotype production was attempted but unsuccessful. **B.** A His-Tag added to the GP C-terminus was used for detection in producer cells and to show incorporation into VSV pseudotypes, except for Tamiami virus, for which incorporation was shown by assaying pseudotype infectivity. GAPDH and VSV-M were used as loading controls for producer cells and viral pseudotypes, respectively.

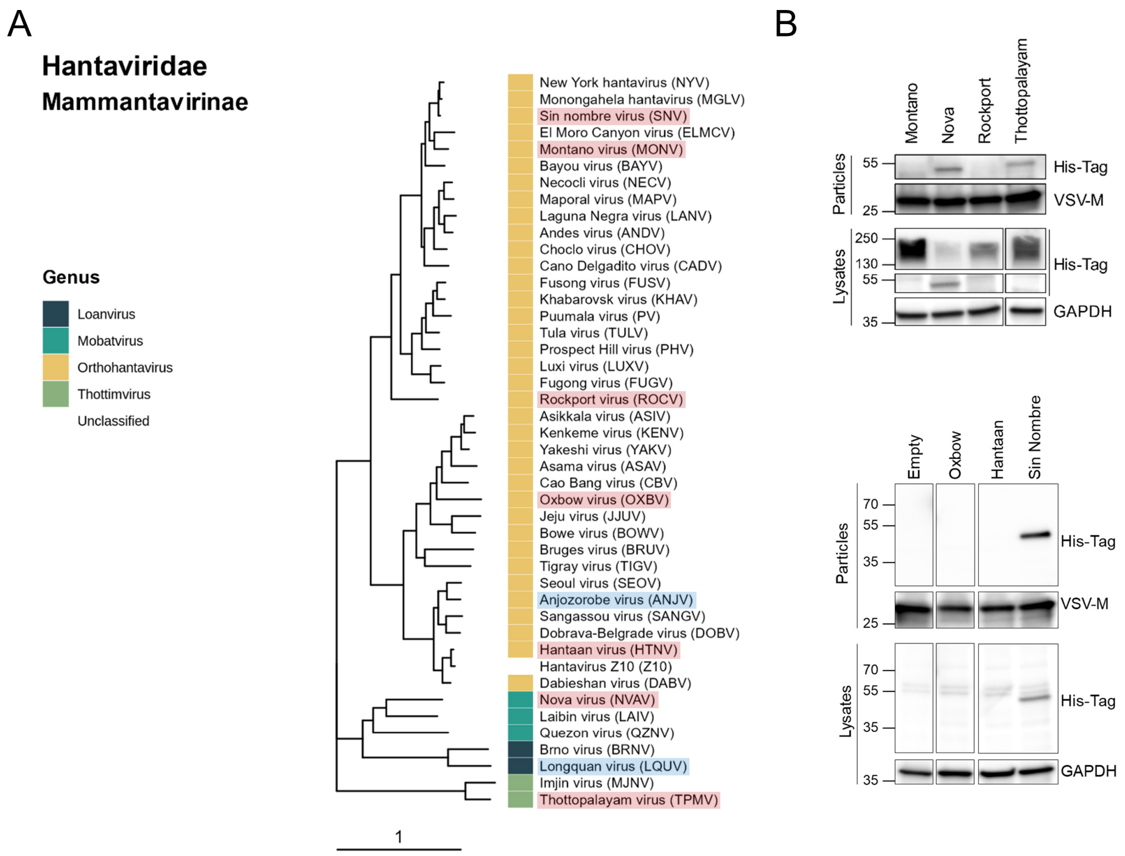


**Figure S8. Mammahantavirinae subfamily Gc-Gn (GPC) phylogeny and Western Blot confirmation of correct pseudotype production. A.** Maximum-likelihood phylogenetic tree, which includes the polyprotein M segment. Species included in our analysis are shaded in red. Species shaded in blue are viruses for which pseudotype production was attempted but unsuccessful. **B.** A His-Tag added to the GPC C-terminus was used for detection in producer cells and to show incorporation into VSV pseudotypes. GAPDH and VSV-M were used as loading controls for producer cells and viral pseudotypes, respectively.

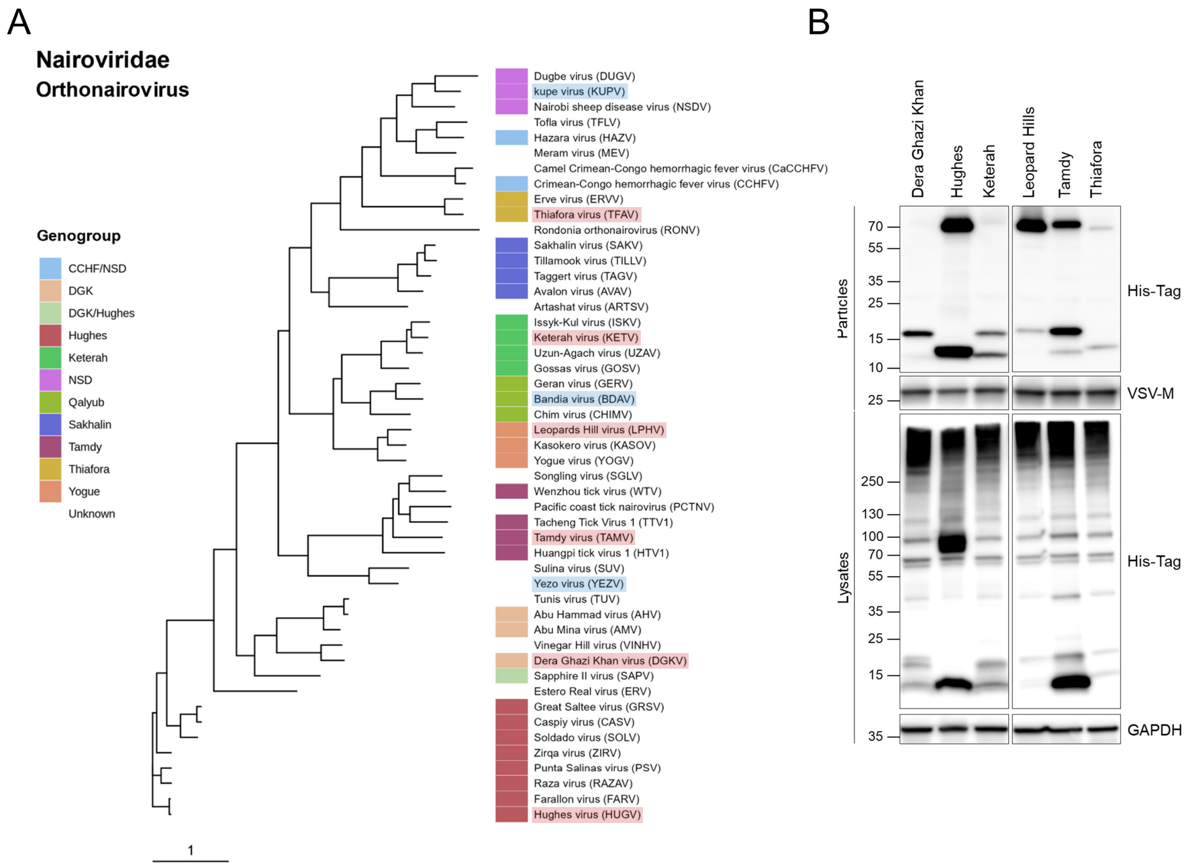


**Figure S9. Nairoviridae Gc-Gn (GPC) phylogeny for the Orthonairovirus genus and Western Blot confirmation of correct pseudotype production. A.** Maximum-likelihood phylogenetic tree, which includes the polyprotein M segment. Species included in our analysis are shaded in red. Species shaded in blue are viruses for which pseudotype production was attempted but unsuccessful. **B.** A His-Tag added to the GPC C-terminus was used for detection in producer cells and to show incorporation into VSV pseudotypes. GAPDH and VSV-M were used as loading controls for producer cells and viral pseudotypes, respectively.

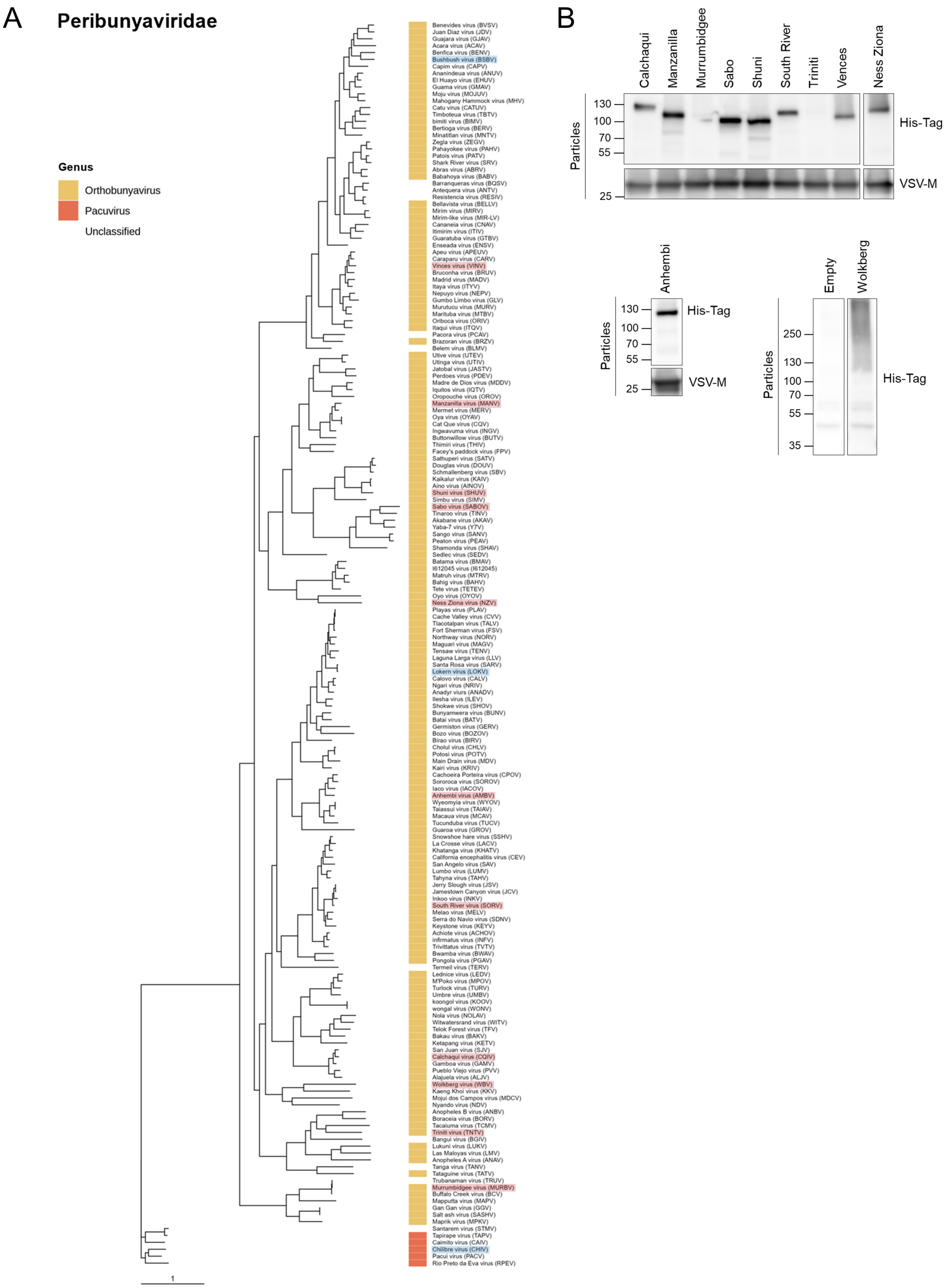


**Figure S10. Peribunyaviridae Gc-Gn (GPC) phylogeny and Western Blot confirmation of correct pseudotype production. A.** Maximum-likelihood phylogenetic tree, which includes the polyprotein M segment. Species included in our analysis are shaded in red. Species shaded in blue are viruses for which pseudotype production was attempted but unsuccessful. **B.** A His-Tag added to the GPC C-terminus was used to show incorporation into VSV pseudotypes. VSV-M was used as a loading control.

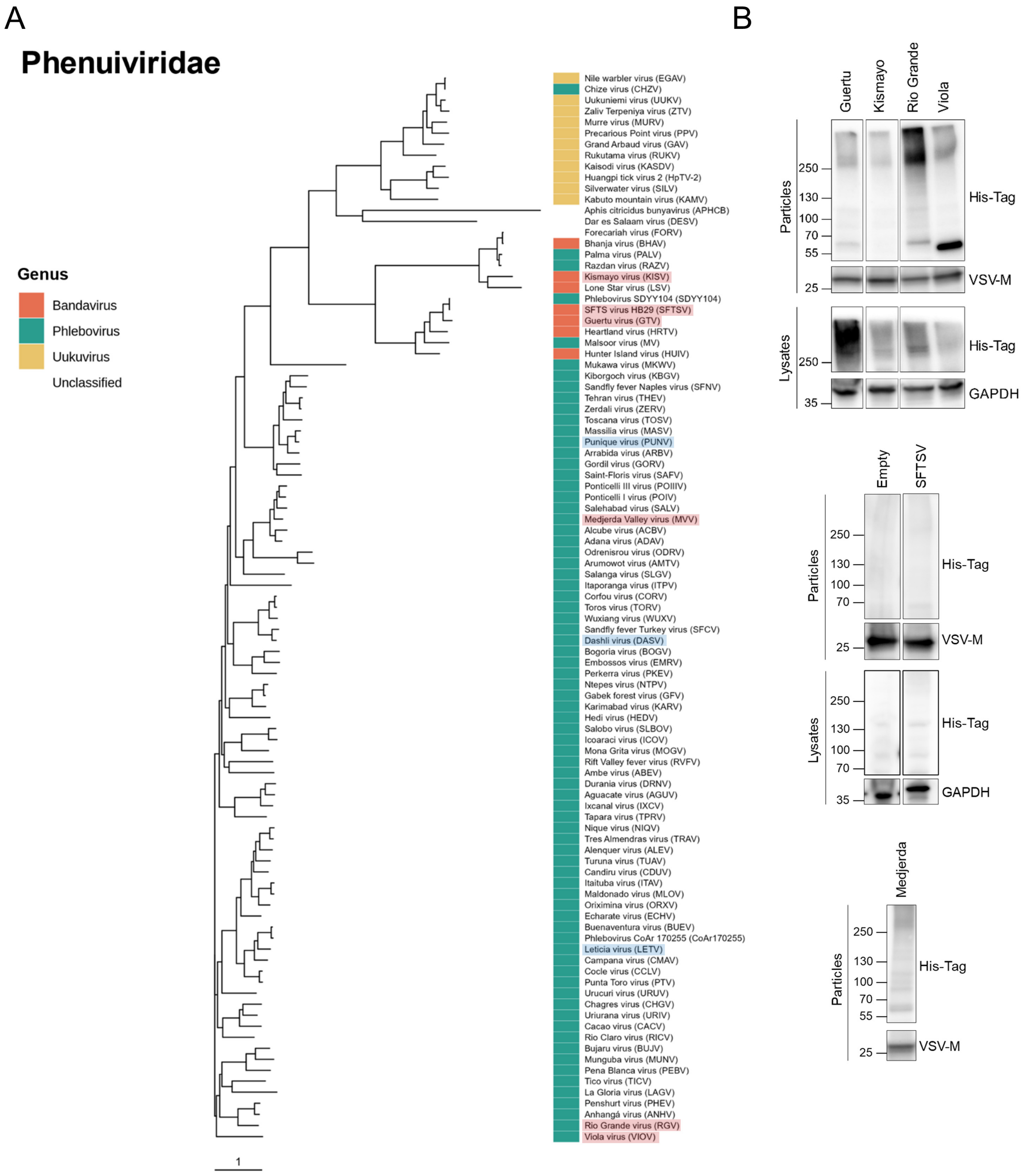


**Figure S11. Phenuiviridae Gc-Gn (GPC) phylogeny and Western Blot confirmation of correct pseudotype production. A.** Maximum-likelihood phylogenetic tree, which includes the polyprotein M segment. Species included in our analysis are shaded in red. Species shaded in blue are viruses for which pseudotype production was attempted but unsuccessful. **B.** A His-Tag added to the GPC C-terminus was used for detection in producer cells and to show incorporation into VSV pseudotypes. GAPDH and VSV-M were used as loading controls for producer cells and viral pseudotypes, respectively.

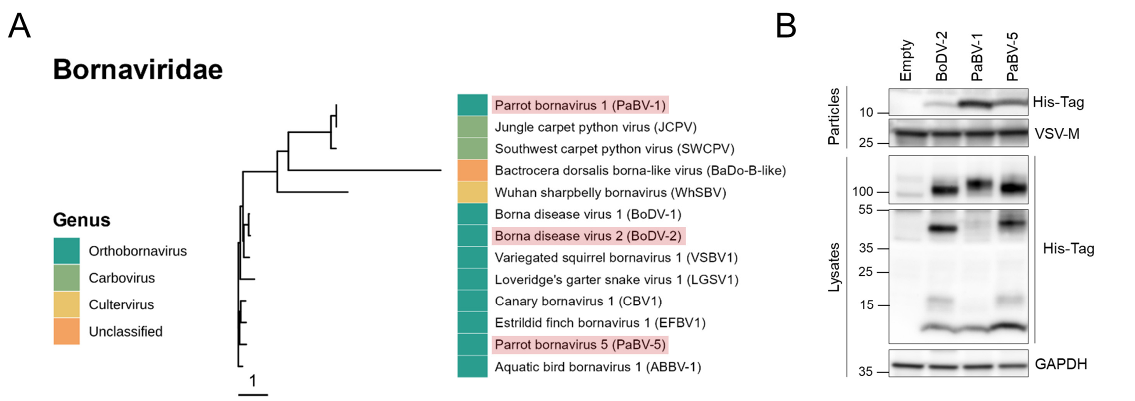


**Figure S12. Bornaviridae G protein phylogeny and Western Blot confirmation of correct pseudotype production. A.** Maximum-likelihood phylogenetic tree. Species included in our analysis are shaded in red. **B.** A His-Tag added to the G protein C-terminus was used for detection in producer cells and to show incorporation into VSV pseudotypes. GAPDH and VSV-M were used as loading controls for producer cells and viral pseudotypes, respectively.

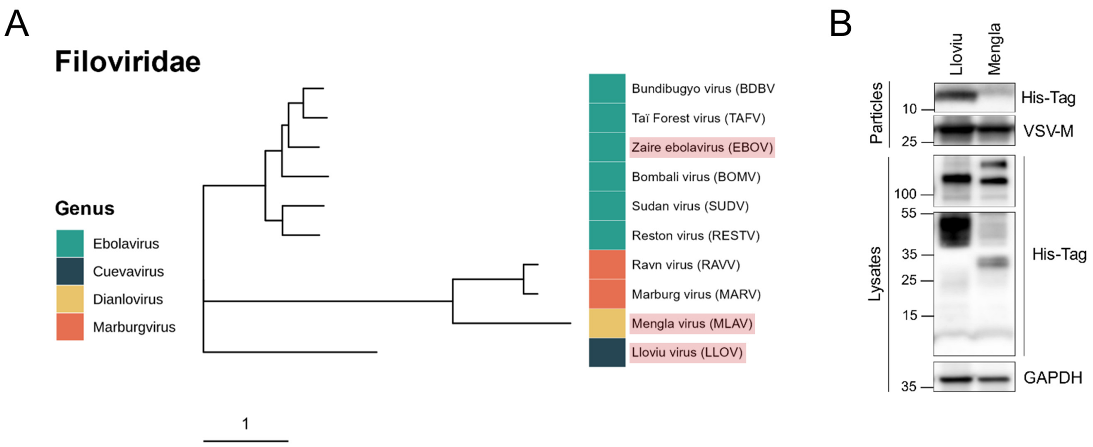


**Figure S13. Filoviridae G protein phylogeny and Western Blot confirmation of correct pseudotype production. A.** Maximum-likelihood phylogenetic tree. Species included in our analysis are shaded in red. **B.** A His-Tag added to the G protein C-terminus was used for detection in producer cells and to show incorporation into VSV pseudotypes, except for Zaire ebolavirus, for which incorporation was shown by assaying pseudotype infectivity. GAPDH and VSV-M were used as loading controls for producer cells and viral pseudotypes, respectively.

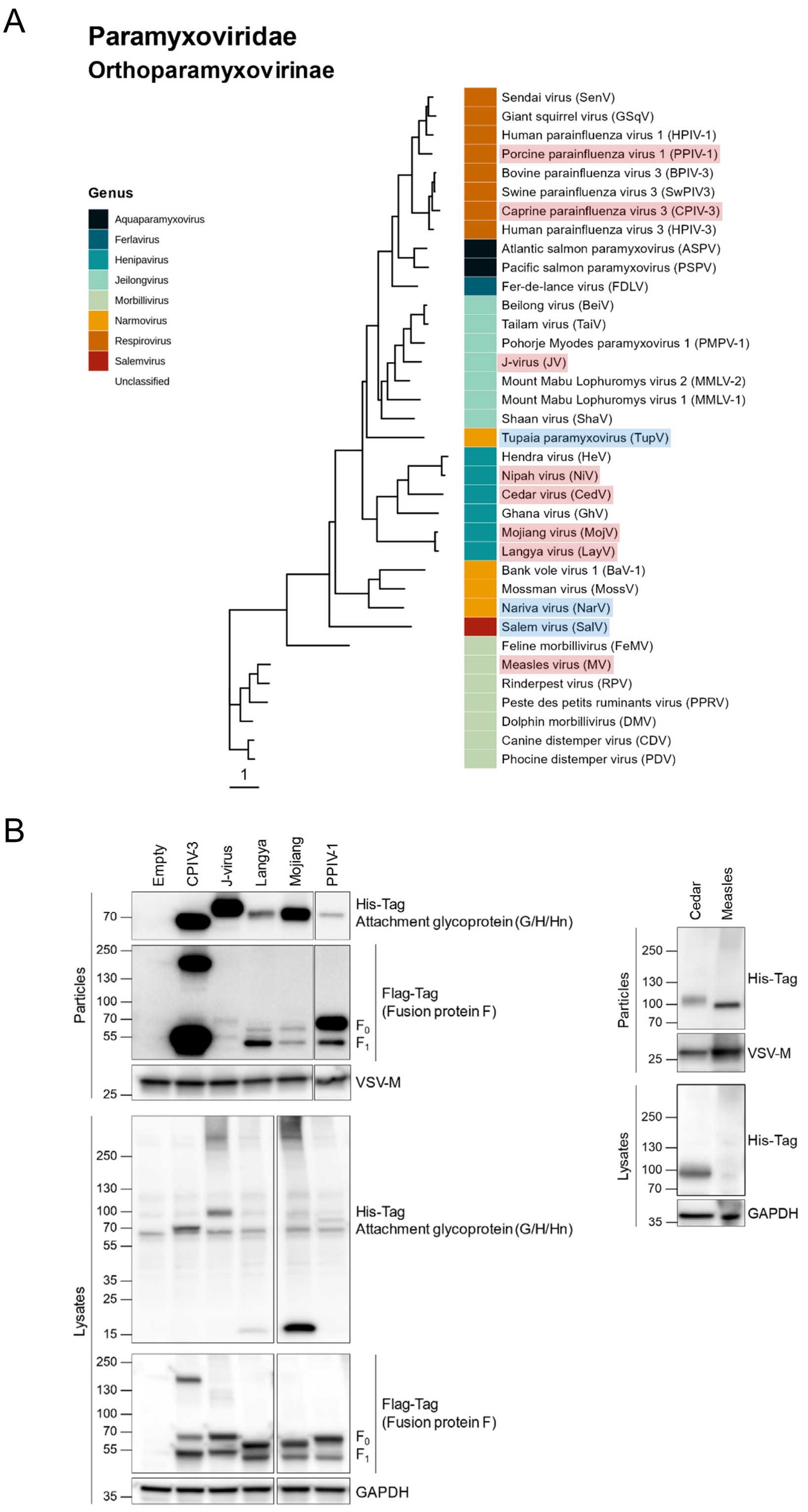


**Figure S14. Orthoparamyxovirinae subfamily attachment glycoprotein (G/H/Hn) and fusion protein (F) phylogeny and Western Blot confirmation of correct pseudotype production. A.** Maximum-likelihood phylogenetic tree of the protein concatenate. Species included in our analysis are shaded in red. Species shaded in blue are viruses for which pseudotype production was attempted but unsuccessful. **B.** A His-Tag added to the G/H/Hn protein N-terminus and a Flag-Tag added to the F protein C-terminus were used for detection in producer cells and to show incorporation into VSV pseudotypes, except for Nipah virus, for which incorporation was shown by assaying pseudotype infectivity. GAPDH and VSV-M were used as loading controls for producer cells and viral pseudotypes, respectively.

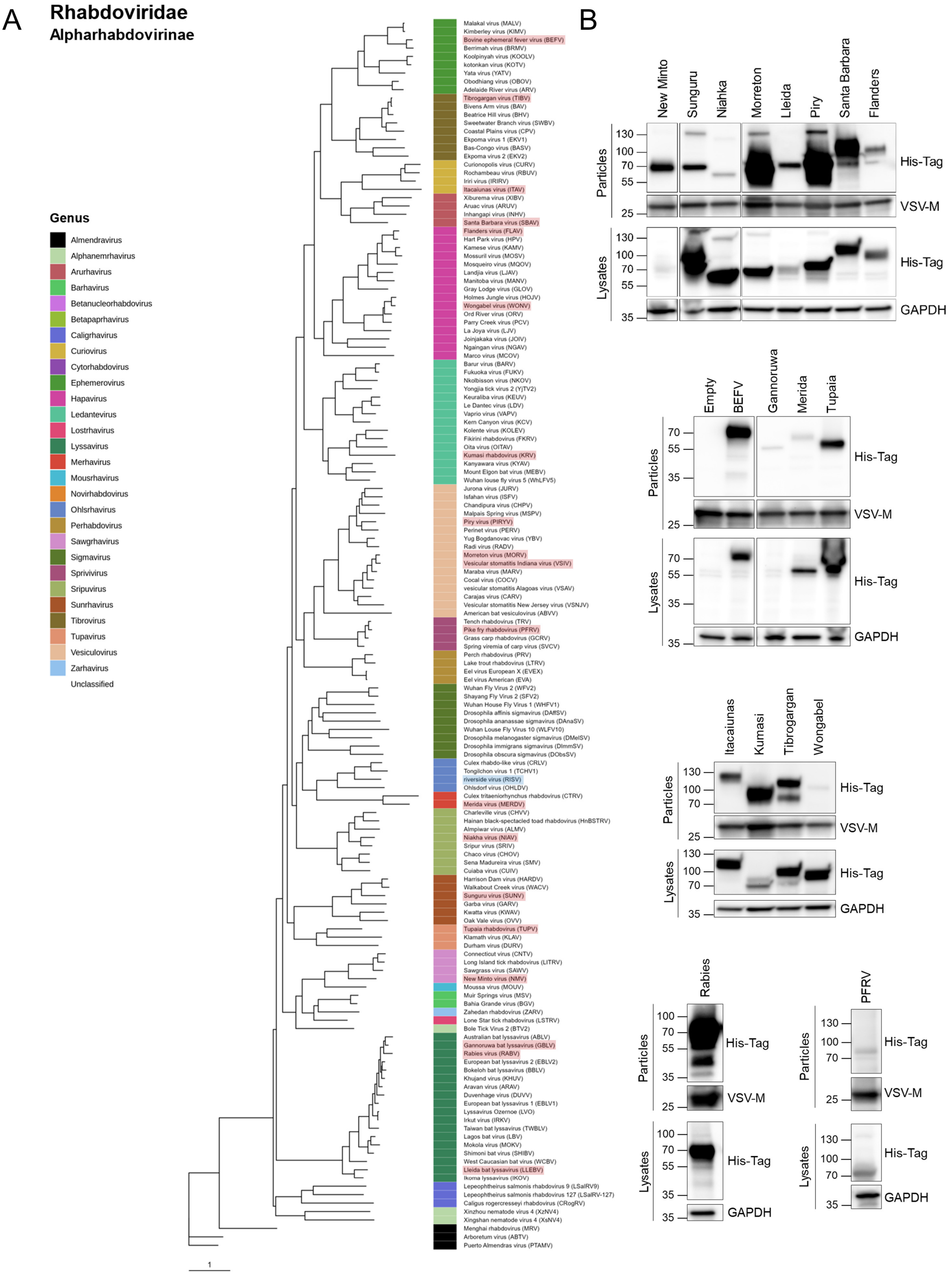


**Figure S15. Apharhabdovirinae subfamily G glycoprotein phylogeny and Western Blot confirmation of correct pseudotype production. A.** Maximum-likelihood phylogenetic tree. Species included in our analysis are shaded in red. Species shaded in blue are viruses for which pseudotype production was attempted but unsuccessful. **B.** A His-Tag added to G protein C-terminus was used for detection in producer cells and to show incorporation into VSV pseudotypes, except for Vesicular stomatitis virus, for which incorporation was shown by assaying pseudotype infectivity. GAPDH and VSV-M were used as loading controls for producer cells and viral pseudotypes, respectively.

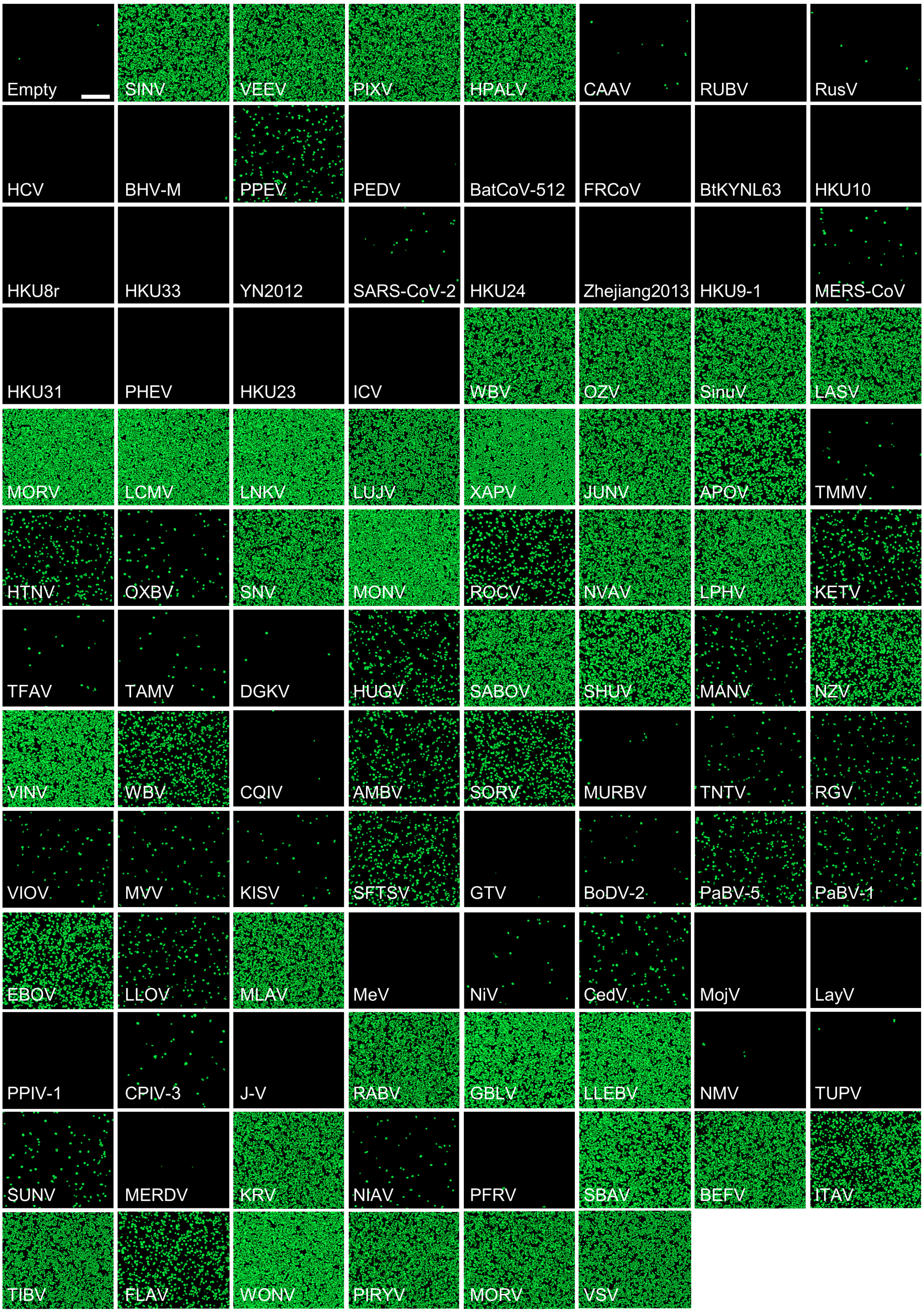


**Figure S16. Preliminary assay of viral pseudotype infectivity in HEK293T cells.** HEK293T cells were infected with viral pseudotypes and imaged 24 hours later with the SX5 Live-Cell Analysis System at a 4X magnification. Images show the GFP-positive area defined using the Incucyte Analysis software. Virus abbreviations are shown. Scale bar: 400 µm.

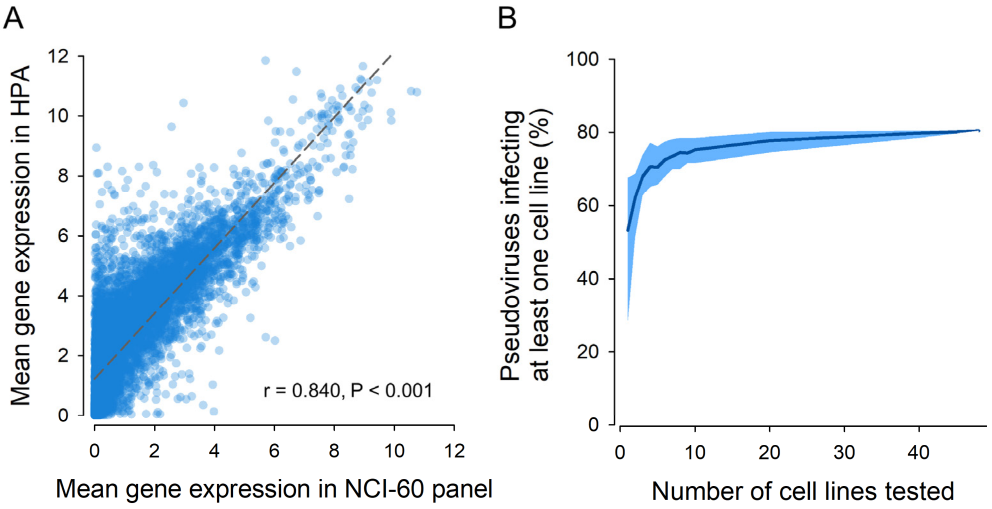


**Figure S17. Suitability of the 48 cells lines for investigating human cell entry. A.** Correlation between the mean mRNA expression values of cell surface genes in the NCI-60 and Human Protein Atlas (HPA) datasets. The expression values in cell lines from the NCI-60 panel used in this study were compared to those of 41 different human tissues available from HPA. Data were retrieved from the corresponding websites, filtered to consider only 7496 annotated as cell surface in Gene Ontology, and expressed as $log_2$(rpkm+1) and $log_2$(nTPM+1), respectively. Pearson r coefficient and p-value are indicated. **B.** Effect of the number of cells tested on estimated human cell infectivity. For each pseudotype, a cell line was defined as infected based on predefined threshold criteria. The number of pseudotypes meeting this condition in at least one cell line was calculated as a function of the number of cell lines considered, from 1 to 48. The dark blue line indicates the average of all possible cell line combinations and the lighter blue shade indicates the range corresponding to the $5^{th}$ and $95^{th}$ percentiles. The percentage of pseudotypes infecting at least one cell line rapidly plateaued, indicating that 48 cell lines is a sufficient sample size to reliably evaluate human cell infectivity.

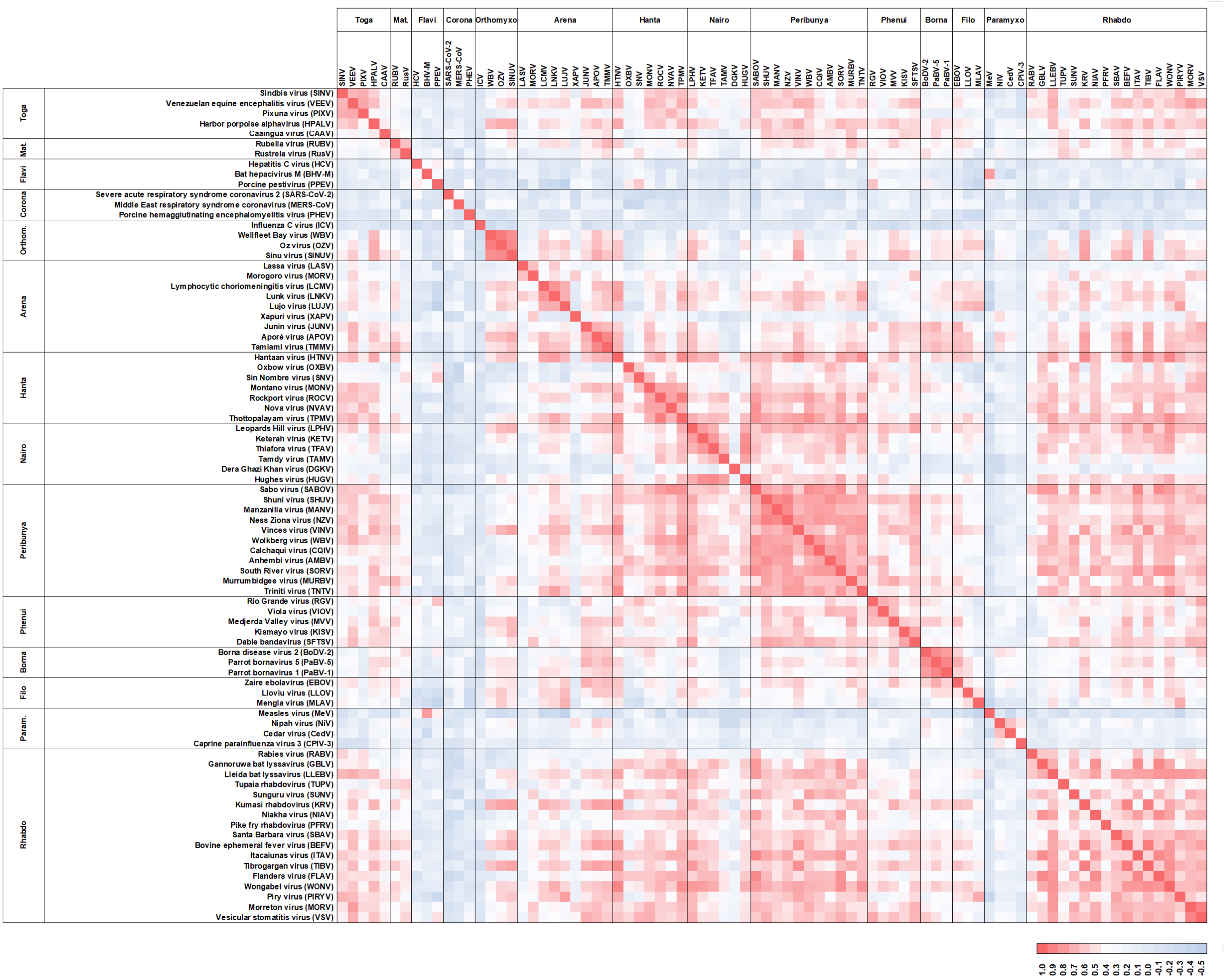

**Figure S18. Correlation matrix between pseudotype infectivity across cell lines.** The heat map indicates the Pearson correlation between each pair of pseudotypes in terms of their $\log_2(R+1)$ values across the 48 cell lines.

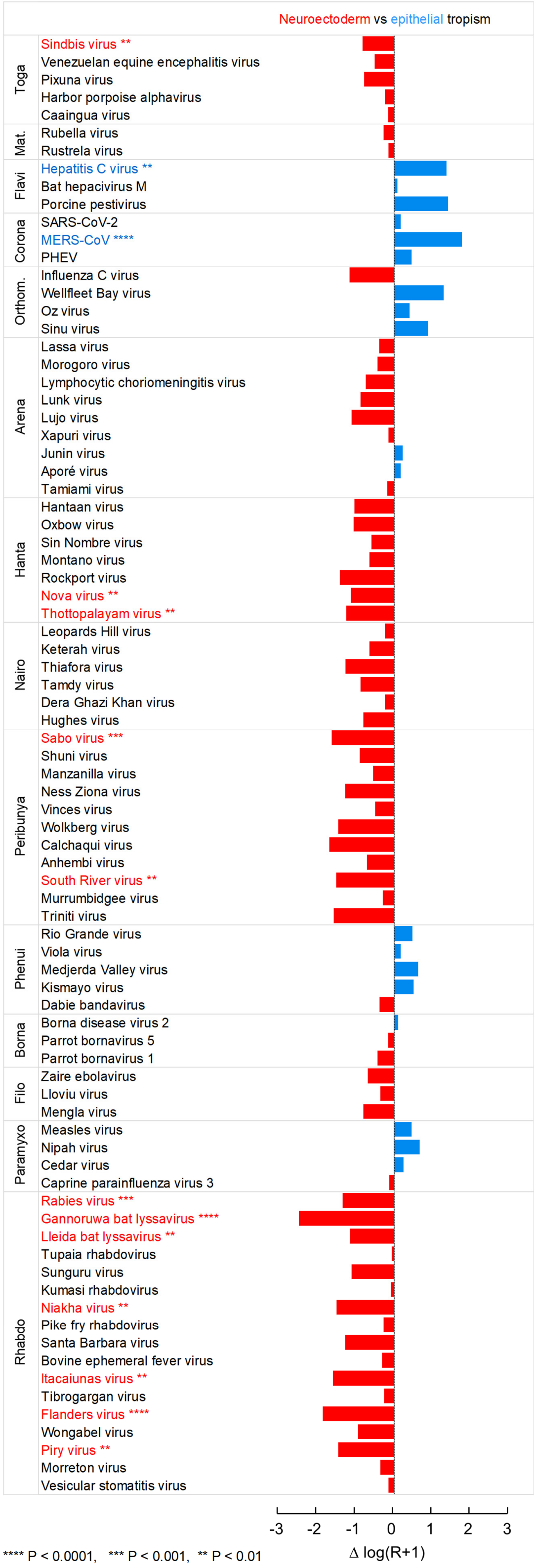


**Figure S19. Relative tropism of pseudotypes for neuroectoderm-derived versus epithelial cells.** For each virus, the difference between the average $\log_2(R+1)$ values in the 15 cell lines derived from the neuroectoderm versus the 33 epithelial cell lines is shown. Red and blue bars show viruses with a preference for neural crest-derived cells and epithelial cells, respectively. The names of the viruses for which this difference was statistically significant according to a Mann-Whitney test are colored accordingly, and significance levels are indicated.

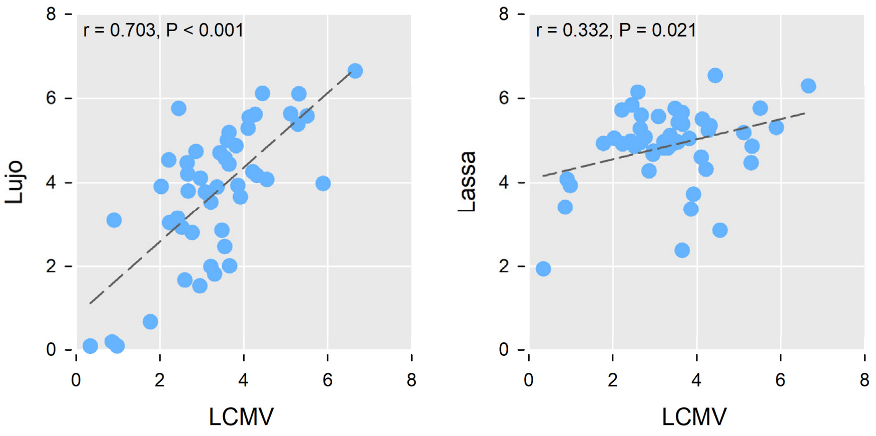


**Figure S20. Infectivity correlation between three arenaviruses.** Left: Lujo versus LCMV pseudotypes. Right: Lassa versus LCMV pseudotypes. Infectivity was measured as $log_2(R+1)$ across the n = 48 cell lines. Each dot represents a cell line. Pearson r coefficients and p-values are indicated.

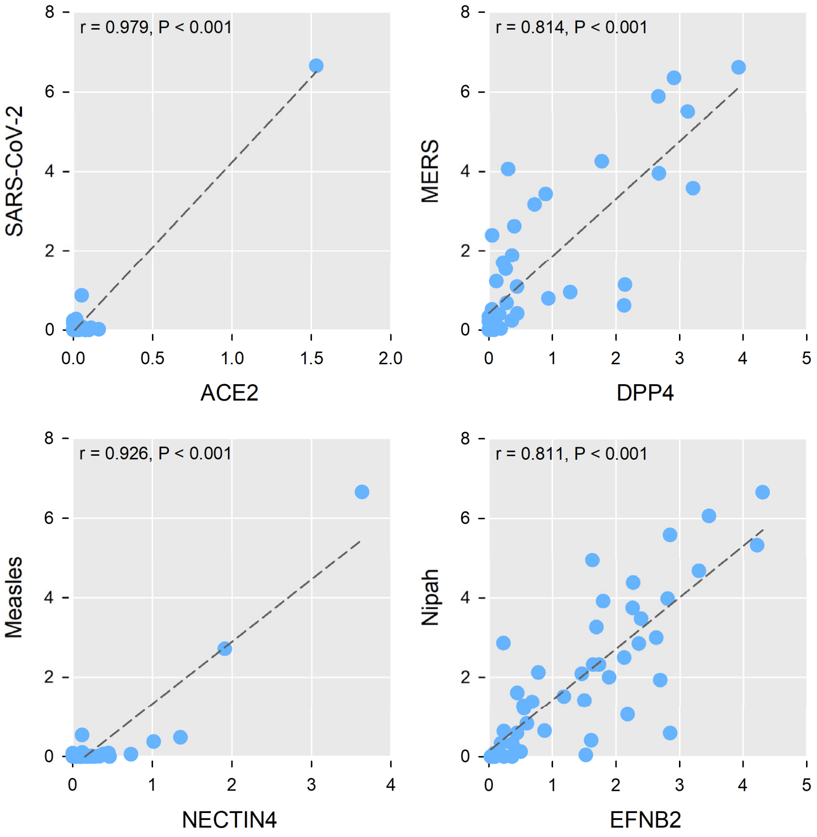


**Figure S21. Examples of significantly positive correlations between the expression of known receptors and pseudotyped virus infectivity.** Correlation between the expression of receptor genes, measured by RNA-seq as $\log_2$(rpkm+1), and viral pseudotypes infectivity, measured as $\log_2$(R+1) across n = 47 cell lines. Each dot represents a cell line. Pearson r coefficients and p-values are indicated.

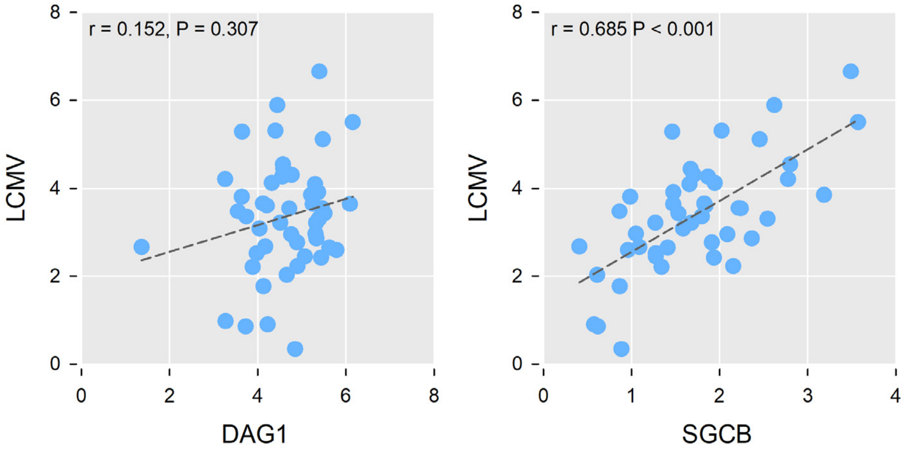


**Figure S22. Example of possible indirect determinants of viral tropism.** LCMV pseudotype infectivity correlated weakly with the mRNA expression levels of the known DAG1 receptor but correlated significantly with those of SCGB, another component of the dystroglycan complex. Each dot represents a cell line (n = 47). Pearson r coefficients and p-values are indicated.

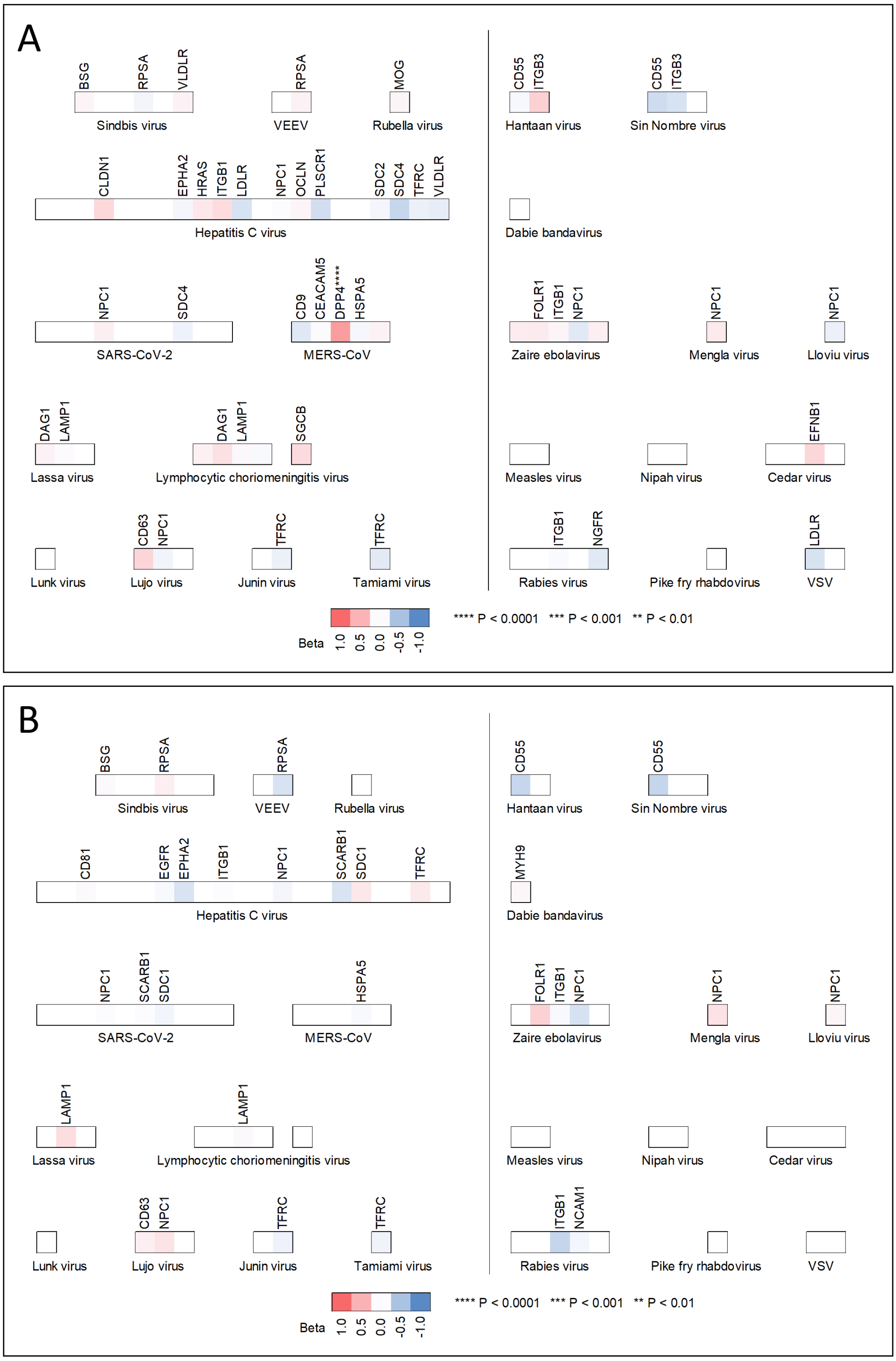


**Figure S23. Analysis of the non-limiting effect of known receptors on viral entry using proteomic data.** Multiple regression analysis of pseudotype infectivity as a function of the protein expression levels of known viral receptors quantified using mass spectrometry. These analyses were performed using two independent quantitative proteomic datasets that included 28 (**A**) and 17 (**B**) receptors[35,36]. Viruses and receptors are shown as in Figure 4, but only for receptors with available proteomic data. The rest of combinations are left blank. Colors indicate the standardized regression coefficient. The dependent variable was $\log_2(R+1)$ in each of the cell lines, and the independent variables were the protein expression levels measured as log-transformed label free quantification intensities.

**Table S1. Features of the RBPs analyzed in this study (Excel format).** The viral family, virus name, RBP name, accession number, fusion protein class, protein size, N-glycosylation sites scaled to protein size, range of the amino acids used for RBP reconstruction, tag used for protein detection, and information about the host range of the virus are provided.

**Table S2. Features of the 48 cell lines of the NCI-60 panel (Excel format).** The name of each cell line, tissue of origin, and age and sex of donor are provided. Additional details are available at dtp.cancer.gov/discovery_development/nci-60/cell_list.htm.

**Table S3. Pseudotype infection data (Excel format).** For each cell-RBP combination, the average log2(R+1) value and a dichotomous variable indicating the presence or absence of infection are provided. Details on the calculation of these variables are provided in the Methods section.

**Table S4. Primers used for cDNA amplification (Excel format).** The gene, accession number, cloning method and vector, and primer sequences are provided.
